# Fiber tractography bundle segmentation depends on scanner effects, vendor effects, acquisition resolution, diffusion sampling scheme, diffusion sensitization, and bundle segmentation workflow

**DOI:** 10.1101/2021.03.17.435872

**Authors:** Kurt G Schilling, Chantal MW Tax, Francois Rheault, Colin B Hansen, Qi Yang, Fang-Cheng Yeh, Leon Y Cai, Adam W Anderson, Bennett A Landman

## Abstract

When investigating connectivity and microstructure of white matter pathways of the brain using diffusion tractography bundle segmentation, it is important to understand potential confounds and sources of variation in the process. While cross-scanner and cross-protocol effects on diffusion microstructure measures are well described (in particular fractional anisotropy and mean diffusivity), it is unknown how potential sources of variation effect bundle segmentation results, which features of the bundle are most affected, where variability occurs, nor how these sources of variation depend upon the method used to reconstruct and segment bundles. In this study, we investigate six potential sources of variation, or confounds, for bundle segmentation: variation (1) across scan repeats, (2) across scanners, (3) across vendors (4) across acquisition resolution, (5) across diffusion schemes, and (6) across diffusion sensitization. We employ four different bundle segmentation workflows on two benchmark multi-subject cross-scanner and cross-protocol databases, and investigate reproducibility and biases in volume overlap, shape geometry features of fiber pathways, and microstructure features within the pathways. We find that the effects of acquisition protocol, in particular acquisition resolution, result in the lowest reproducibility of tractography and largest variation of features, followed by vendor-effects, scanner-effects, and finally diffusion scheme and b-value effects which had similar reproducibility as scan-rescan variation. However, confounds varied both across pathways and across segmentation workflows, with some bundle segmentation workflows more (or less) robust to sources of variation. Despite variability, bundle dissection is consistently able to recover the same location of pathways in the deep white matter, with variation at the gray matter/ white matter interface. Next, we show that differences due to the choice of bundle segmentation workflows are larger than any other studied confound, with low-to-moderate overlap of the same intended pathway when segmented using different methods. Finally, quantifying microstructure features within a pathway, we show that tractography adds variability over-and-above that which exists due to noise, scanner effects, and acquisition effects. Overall, these confounds need to be considered when harmonizing diffusion datasets, interpreting or combining data across sites, and when attempting to understand the successes and limitations of different methodologies in the design and development of new tractography or bundle segmentation methods.

## Introduction

Diffusion-weighted magnetic resonance imaging (dMRI) has proven valuable to characterize tissue microstructure in health and disease [1-3]. Moreover, the use of dMRI fiber tractography to virtually dissect fiber pathways [4] is increasingly used to localize microstructure measurements to specific white matter bundles [5-7], and to study the connections and shapes of pathways [4, 8-16]. Despite promises of noninvasive measurements of white matter features, variability may exist in measurements due to inherent variability within scanners and across scanners, differences in acquisition protocol parameters, and differences due to processing pipelines, amongst others. These sources of variance challenge the quantitative nature of derived measures of microstructure and connectivity, and hinder the ability to interpret different findings or combine different datasets.

These effects have been intensively studied for tissue microstructure features, specifically diffusion tensor imaging (DTI)[17] indices of fractional anisotropy (FA) and mean diffusivity (MD). Numerous studies have characterized intra-scanner and inter-scanner DTI variability [18-29], differences due to acquisition parameters [24, 25, 28, 30-33] including image resolution, number of diffusion images, and diffusion sensitization (i.e. the b-value), and differences due to processing and algorithmic choices [34, 35]. These have paved the way towards recommendations and guidelines for reliable and reproducible DTI [36-39]; however, a standardized universal dMRI protocol does not exist, and differences are expected across sites and studies (Figure 1) [40, 41]. Yet, there is significant interest in combining data from different sites to increase statistical power and benefit from multi-center recruitment abilities [19, 40, 42-48], and it is clear that these differences need to be accounted for, or removed, prior to data aggregation or joint statistical analysis.

**Figure 1.**
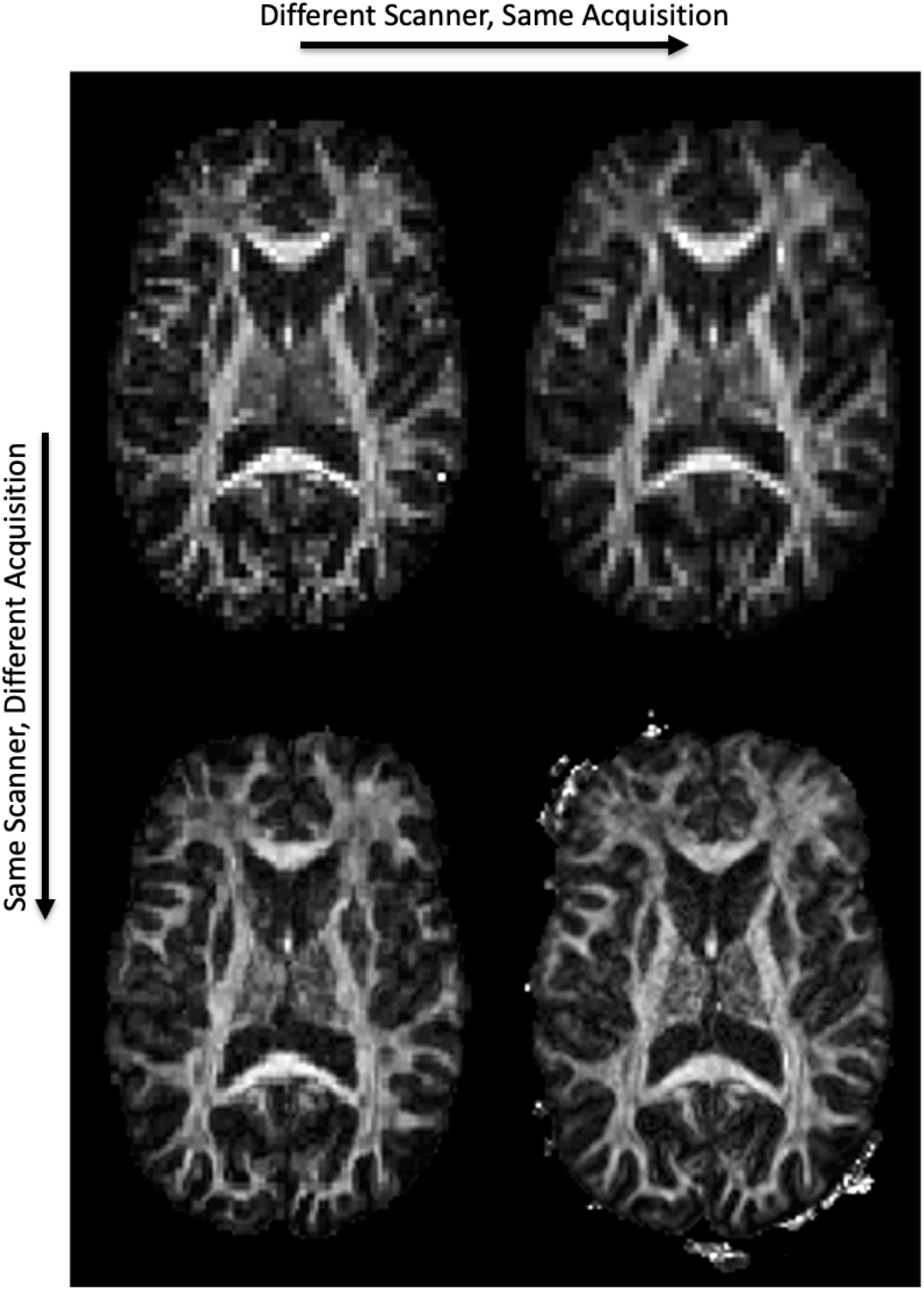
Microstructure varies across scanners and across acquisitions. An FA map is shown, derived from the same subject, on two scanners (Siemens Prisma, left; Siemens Connectom, right) and two acquisitions (standard acquisition, top; state-of-the-art acquisition, bottom). See Methods for scanner and acquisition details.

Notwithstanding the increased awareness and improved characterization of dMRI microstructural measures, very little work has been performed to characterize and understand reproducibility of tractography-derived features across scanners, across protocols, and across different tractography bundle segmentation algorithms [49, 50]. Variability in tractography estimates of fiber pathways will further increase variability in connectivity analyses and impact microstructural characterization, e.g. when tractography is used to define ROIs or to perform along-tract profiling. While few studies do exist, they are often limited to a single pathway [51, 52], a single dissection protocol [53, 54], or a single source of potential variation [55], such as test-retest or population-based reproducibility [54, 56, 57]. Additionally, they do not investigate where in the brain or along the pathway that this variability occurs, and are often limited to characterizing only microstructural features of these pathways (i.e., the FA or MD along or within the pathway) [26, 58]. Thus, we currently do not which sources of variation impact tractography bundle segmentation the most, which features of the bundle are most affected, where variability occurs, nor how these questions are dependent upon the workflow used to dissect fiber bundles. Thus, for the first time, we combine, assess, and rank all previously studied sources of potential variation in the same study, with a focus on tractography rather than just DTI measures.

Here, we investigate and compare the reproducibility of tractography across six confounds, or sources of variation: intrinsic variability across scan repeats, differences across scanners, across vendors, across different acquisition spatial resolution and acquisition angular resolution, and across different diffusion sensitizations (b-values). We employ and examine four fully-automated and commonly utilized bundle reconstruction workflows on two cross-scanner cross-protocol benchmark datasets. We first investigate how these confounds affect not only the overlap and location of pathways, but also evaluate variability in topological measures of the bundle including length, area, shape, and volume features. We ask which pathways, which bundle segmentation workflow, and which features are most reproducible? And what source of variation is most significant for each method? Second, we visualize *where in the brain, and where within a pathway*, tractography is most variable (and most robust) and investigate if sources of variation effect this in different ways. Third, we quantify and visualize differences in tractography that result when using different bundle segmentation workflows. Finally, we analyze traditional DTI measures and quantify differences due to these sources of variation as well as the *added* variance introduced by the tractography process over and above that inherent across scanners and across acquisition protocols.

## Methods

### Datasets

Here we utilize two open-sourced multi-subject, multi-scanner, and multi-protocol benchmark databases: the MASiVar [59] and MUSHAC datasets [40, 41]. We note that other multi-site databases exist (see Discussion), although they are often limited to investigating differences across subjects and scanners, whereas the two chosen datasets together allow investigation of repeats, scanners, vendors, and acquisition protocols (resolution, direction, b-values).

#### MUSHAC dataset

The MUSHAC database will allow investigation of cross-scanner, cross-protocol, and cross b-value effects [40, 41]. This database was part of the 2018 and 2019 MICCAI Harmonization challenge. Here, we utilize the data acquired from 10 healthy subjects used as training data in the challenge, and described in [40, 41]. Each subject has 4 unique datasets. This work focuses on the data acquired on two scanners with different gradient strengths: a) 3T Siemens Prisma (80 mT/m), and b) 3T Siemens Connectom (300 mT/m). Two types of protocols were acquired from each scanner: 1) a ‘standard’ protocol with acquisition parameters matched to a typical clinical protocol; and 2) a more advanced or ‘state-of-the-art’ protocol where the superior hardware and software specifications were utilized to increase the number of acquisitions and spatial resolution per unit time. The ‘standard’ protocol from both scanners is matched as closely as possible, with an isotropic resolution of 2.4 mm, TE=89 ms and TR=7.2 s, and 30 diffusion-weighted directions acquired at two b-values: b = 1200, 3000 s/mm2 (scan time ∼7.5 minutes). On the other hand, the Prisma ‘state-of-the-art’ data has a higher isotropic resolution of 1.5 mm, TE=80 ms, TR=7.1s and 60 directions at the same b-values (∼14.5 minutes). While the Connectom ‘state-of-the-art’ data has the highest resolution of 1.2 mm with TE=68 ms, TR=5.4 s and 60 directions (∼11 minutes). All data was corrected for distortions, motion, eddy currents [60], and gradient nonlinearity distortions [61]. For each subject, the Prisma standard-acquisition dataset was used as a reference space and all additional datasets were affinely registered to this space using the corresponding FA maps with FSL Flirt with appropriate b-vector rotation.

#### MASiVar dataset

The MASiVar database will allow investigation of scan-rescan and cross-scanner effects. Here we used a subset of Cohort II of this database described in [59], which consisted of 5 healthy subjects with 6 unique “datasets”. Each subject was scanned on four scanners: a) 3T Philips Achieva (80 mT/m) and b) a different 3T Philips Achieva (60mT/m) at the same site, c) a 3T General Electric Discovery MR750 Scanner at a different site, and d) a 3T Siemens Skyra scanner at a different site. These acquisitions were matched as closely as possible and are similar to that of the standard-protocol described above: with an isotropic resolution of 2.5 mm, TE=55 ms and TR=6.2s (7.0s for scanner-b), and 32 diffusion-weighted directions acquired at b = 1000 s/mm2 (scan time ∼3.5 minutes). Additionally, the subjects were scanned twice on the first scanner, and also had an acquisition that consisted of a 96-direction b=1000 dataset, both of which were also utilized in the current study. We note that one subject did not have a repeat scan on the first scanner (a) and one subject did not have a scan on the GE Scanner (b).

All data were corrected for distortions, motion, and eddy currents [60, 62]. For each subject, the first session on scanner-a was used as a reference space and all additional datasets were affinely registered to this space using the corresponding FA maps with FSL Flirt [63] with appropriate b-vector rotation.

### Sources of variation

We investigate several possible sources of variation in the bundle segmentation process.

***RESCAN: the effects of repeating a scan on the same scanner (i***.***e. scan-rescan) in a different session, but with a matched acquisition. This effect is quantified using the repeated acquisitions from the MASiVar database***.

***SCAN1: inter-scanner (cross-scanner) effects, with a matched acquisition and of the same vendor. SCAN1 is quantified using the matched acquisitions from the MASiVar database acquired on different Philips scanners (both Philips Achieva)***.

***SCAN2: inter-scanner (cross-scanner) effects, with a matched acquisition and of the same vendor. SCAN2 is quantified using the matched standard acquisitions from the MUSHAC acquired on different Siemens scanners (Siemens Connectom and Siemens Prisma)***.

***VEN1: inter-vendor (cross-vendor) effects, with a matched acquisition. VEN1 is quantified using the matched acquisitions from the MASiVar database, but acquired on scanners from different vendors (Philips Achieva and General Electric Discovery)***.

***VEN2: inter-vendor (cross-vendor) effects, with a matched acquisition. VEN2 is quantified using the matched acquisitions from the MASiVar database, but acquired on scanners from different vendors (Philips Achieva and Siemens Skyra)***.

***RES1: effects of spatial resolution, with matched scanner, diffusion directions, and b-value. RES1 is quantified by using the MUSHAC acquisitions from the Prisma standard-acquisition and from the Prisma state-of-the-art acquisition but with only 30 uniformly distributed directions utilized (to match the standard-acquisition). This represents differences between a 2***.***4mm isotropic and 1***.***5mm isotropic acquisition***.

***RES2: effects of spatial resolution, with matched scanner, diffusion directions, and b-value. RES2 is quantified by using the MUSHAC acquisitions from the Connectom standard-acquisition and from the Connectom state-of-the-art acquisition but with only 30 uniformly distributed directions utilized (to match the standard-acquisition). This represents differences between a 2***.***4mm isotropic and 1***.***2mm isotropic acquisition***.

***DIR1: effects of number of diffusion-weighted directions, with matched scanner, resolution, and b-value. DIR1 is quantified using the MASIvar acquisitions from the first scanner at 32 directions and the acquisition on the same scanner at 96 directions***.

***DIR2: effects of number of diffusion-weighted directions, with matched scanner, resolution, and b-value. DIR2 is quantified using the MUSHAC acquisitions from the state-of-the art Prisma acquisition with only 30 uniformly distributed directions utilized and the full state-of-the art acquisition which consists of 60 directions***.

***BVAL: effects of changing the b-value, on the MUSHAC Prisma scanner with the ‘standard’ protocol, from b=1200 to b=3000, within the same acquisition***.

We note that we also investigated a second effect of b-value (within the state-of-the art Prisma protocol, with no statistically significant differences, and for figure simplicity only show the above-mentioned b-value analysis). Previous version of this manuscript (and preprint) included an ACQ1 and ACQ2 (from state-of-the-art to standard-acquisition) that were isolated into both effects of directions (DIR1 and DIR2) and resolution (RES1 and RES2).

A final source of variation investigated is that caused by the use of different bundle reconstruction workflows. Because all workflows segment different numbers of, and sets of, fiber pathways (see below), for this analysis, we investigated only those fiber pathways which are common to all algorithms. In this case, we identified 7 (bilateral) pathways which are segmented by all automated methods.

### Tractography bundle dissection

We utilized four common, well-validated, and fully-automated fiber bundle reconstruction workflows, all implemented using standard and/or recommended settings. It is important to highlight that each workflow included differences in local fiber-direction estimation, fiber tractography, and bundle segmentation algorithms, and our attempt was to implement the entire workflow as would be done in a typical scientific study (see Discussion on limitations of confounds due to differences in bundle segmentation process). While there are dozens of bundle segmentation algorithms, we have chosen these to be representative of common approaches, utilizing regions of interest, atlases, machine learning, templates, etc. (see Discussion and Limitations).

#### TractSeg

TractSeg is based on convolutional neural networks and performs bundle-specific tractography based on a field of estimated fiber orientations [64-66]. We implemented the dockerized version at (https://github.com/MIC-DKFZ/TractSeg), which generates fiber orientations using constrained spherical deconvolution with the MRtrix3 software [67]. We note that different reconstruction methods could have been chosen to generate fiber orientations. This method dissects 72 bundles.

#### Automatic Fiber Tractography (ATK)

ATK was performed in DSI Studio software using batch automated fiber tracking [68]. Data were reconstructed using generalized q-sampling imaging [69] with a diffusion sampling length ratio of 1.25.

20 white matter pathways were automatically reconstructed using seeding regions defined in the HCP842 tractography atlas [70], randomly generated tracking parameters of anisotropy threshold, angular threshold, step size, and subsequent segmentation and pruning. The Dockerized source code is available at http://dsi-studio.labsolver.org.

#### Recobundles (RECO)

Recobundles [71] segments streamlines based on their shape-similarity to a dictionary of expertly delineated model bundles [70]. Recobundles was run using DIPY [72] software (https://dipy.org) after performing whole-brain tractography using spherical deconvolution and DIPY LocalTracking algorithm. The bundle-dictionary contains 80 bundles, but only 44 were selected to be included in this study after consulting with the algorithm developers based on internal quality assurance (for example, removing cranial nerves which are often not used in brain imaging). Of note, Recobundles is a method to automatically extract and recognize bundles of streamlines using prior bundle models, and the implementation we chose uses the DIPY bundle dictionary [70] for extraction, although others can be used, as well as alternative shape-similarity filtering criteria.

#### Xtract

Xtract (https://fsl.fmrib.ox.ac.uk/fsl/fslwiki/XTRACT) is a recent automated method for probabilistic tractography based on carefully selected inclusion, exclusion, and seed regions, defined in a standard space [73]. Xtract used the ball-and-stick model of diffusion from FSL’s bedpostx algorithm [63], in combination with a probabilistic tractography algorithm probtrackx, to reconstruct 42 white matter pathways. In contrast to the preceding methods, which result in streamlines, this method results in visitation count maps for each pathway.

A list of all segmentations generated from each method and corresponding acronyms is given in the appendix. The 7 pathways identified to be common to all tractography bundle segmentation techniques includes: arcuate fasciculus (AF), corticospinal tract (CST), inferior fronto-occipital fasciculus (IFO), inferior longitudinal fasciculus (ILF), middle longitudinal fasciculus (MdLF), optic radiations (OR), and uncinate fasciculus (UF), all of which are bilateral including left (_L) and right (_R) hemisphere pathways.

A thorough quality control was performed for all subjects, and for all pathways. This included first visualization and verification of adequate alignment of all FA maps (to ensure appropriate quantification of overlap measures). Second, all pathways, for all subjects, were visualized in mosaic form using tools from the SCILPY toolbox (https://github.com/scilus/scilpy), and pathways were visually assessed and removed from analysis if deemed in the incorrect location or shape.

Finally, individual bundles were removed from analysis if the number of segmented streamlines was less than 3 standard deviations away from the mean number (for each pathway), or if the total number of streamlines was below 200 (indicating failure of tractography), and subjects were removed from analysis (for a given algorithm) if >20% of pathways failed QC.

### Feature extraction

A number of features were extracted from each bundle segmented. First, for simple comparisons of the volume occupied by each pathway, all bundles (from all methods) were binarized and resampled at 1mm isotropic resolution. For methods generating streamlines (Tractseg, ATK, and RECO) this is equivalent to binarizing based on a streamline density of 1. Because Xtract output is in the form of a normalized probability distribution, where a threshold of 2.5E-4 was chosen based on [73]. The binarized segmentation was used for measures of Dice overlap (described below).

Second, several descriptors of the shape and geometry of the bundles were extracted. Shape analysis was performed using DSI Studio, and made available as matlab code (https://github.com/dmitrishastin/tractography_shapes/), based on [68], to derive length, area, volume, and shape metrics of a bundle. Briefly, length features include mean length, span, diameter, and average radius of end regions. Area features include total surface area and the total area of end regions. Volume features include total volume, trunk volume, and branch volume. Shape features include pathway curl, elongation, and irregularity.

Finally, microstructure measures of FA and MD (calculated using iteratively reweighted linear least squares estimator) within pathways were extracted. In all cases, a simple measure of the average value within the binary volume was performed, although we note that these measures can also be weighted by certainty or streamline density. To isolate the added variation due to tractography from that of the existing sources of variation, these measures were extracted in two ways. First, using the binary regions defined in the reference scan-space only (i.e., the Prisma standard-acquisition and first session on scanner-a for MUSHAC and MASiVar datasets respectively) were used as the *same* region-of-interest across all effects, in order to isolate each source of variation while keeping ROIs constant. Second, the binary region defined by tractography for each specific dataset was used to extract the average FA (or MD), which includes both variation due to the effect under investigation and the variation due to tractography differences.

### Reproducibility Evaluation

Reproducibility was evaluated using several metrics, and across each source of variation. First, the Dice overlap was calculated for each pair of bundles as an overall measure of similarity of volumes. The Dice overlap is calculated as two times the intersection divided by the sum of the volumes of each dataset. Results were displayed across all fiber pathways for a given source of variation, and differences between effects were calculated using the nonparametric paired (i.e. same subject, different effect) Wilcoxon signed rank tests.

Differences in scalar shape features are calculated as the mean absolute percentage error (MAPE), sometimes referred to as the mean absolute percentage deviation. For two different scans, this measure is calculated as the difference divided by the mean, and can be converted to a percentage error by multiplication by 100. This measure was calculated over all subjects, and results were displayed across all fiber pathways for a given source of variation. Differences between effects were again calculated using the nonparametric paired (i.e. same subject, different effect) Wilcoxon signed rank tests.

For visual comparisons only, all subjects were nonlinearly registered to MNI space, using the 1mm isotropic FA template and the corresponding FA maps with FSL FLIRT + FNIRT. Streamlines were directly warped to this space for visualization of agreement/disagreement across the cohort. Note that quantification of shape features was performed in native space prior to warping.

For all statistical analysis, thresholds were corrected for multiple comparisons. For example, when investigating differences in effects of DICE/MAPE, etc., we tested differences between 10 effects, resulting in 55 tests performed for each analysis.

## Results

### Qualitative Variation

**Figure 1** shows FA maps of the same subject, but acquired on different scanners and with different protocols. In agreement with the literature [40, 41], differences in magnitude, contrast, and signal-to-noise ratios are readily apparent, and dMRI measures qualitatively vary due to scanner and acquisition effects.

**Figure 2** shows tractography bundle segmentation results for an example pathway (the arcuate fasciculus; AF) on a single subject, for two scanners, two protocols, two b-values, and all four reconstruction methods. For a given bundle segmentation method, minor differences are observed in individual gyri and at regions of low streamline density. However, bundles are visually very similar across scanners and protocols, with similar shapes, locations, curvatures, and connections. Most notably, and as expected [55], the biggest differences are observed when comparing the same pathway across different bundle segmentation methods.

**Figure 2.**
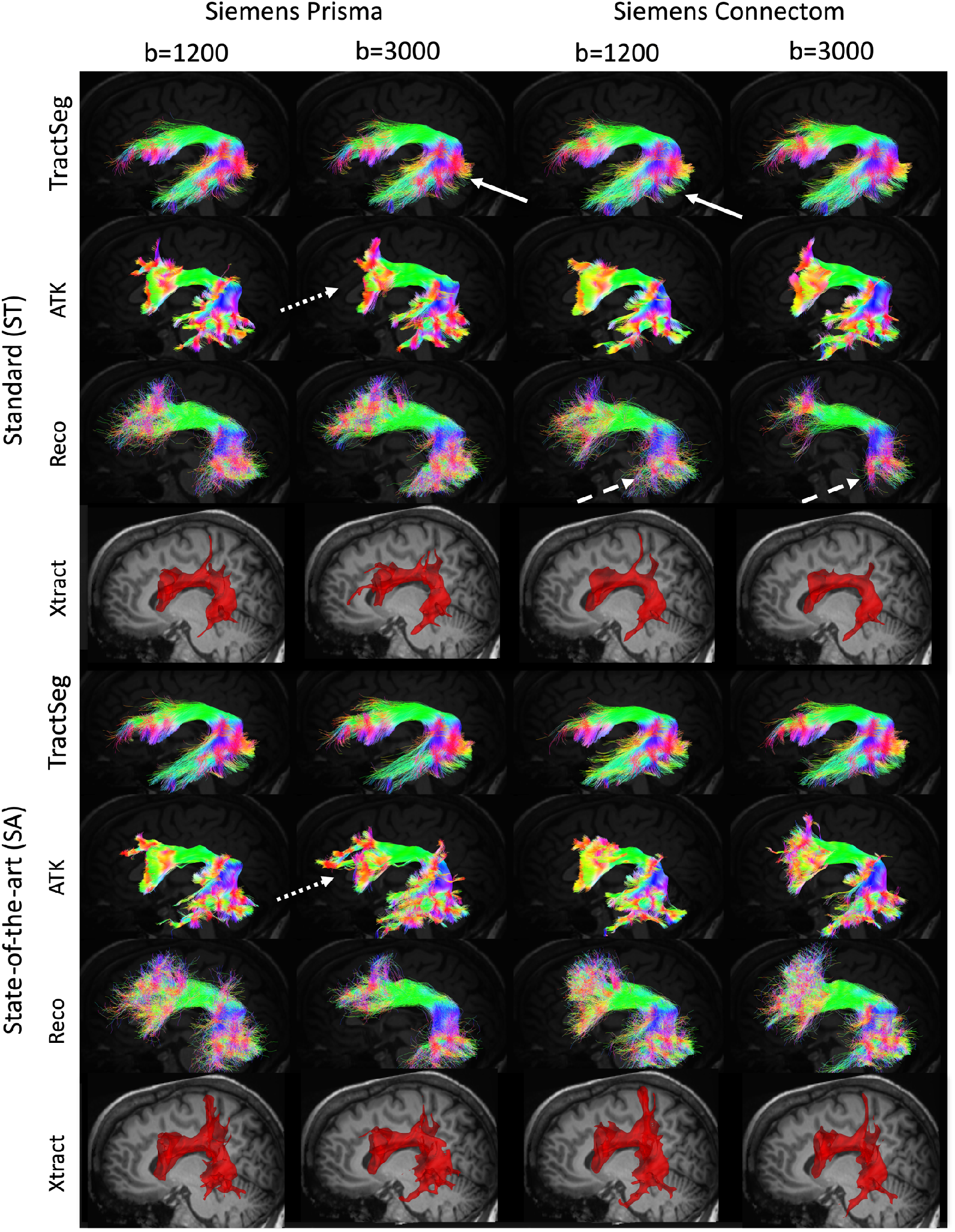
Tractography varies across scanners, acquisitions, b-values, and bundle segmentation methods. On the same subject, the arcuate fasciculus is shown for each of the 4 bundle segmentation methods, for two scanners and two acquisitions. Note that the pathway is visualized as streamlines for TractSeg, ATK, and Reco but a probability density map for Xtract. Arrows highlight visible examples of differences in streamlines across scanners (solid arrows), across acquisition (dotted arrows), and across b-values (dashed arrows).

### Quantitative variation due to rescan, scanner, vendor, resolution, directions, and b-value effects

The effects of RESCAN, SCAN1, SCAN2, VEN1, VEN2, DIR1, DIR2, and BVAL on Dice overlap coefficient is shown in **Figure 3** for fourteen selected pathways common to all bundle segmentation methods. Notably, reproducibility is most dependent on the bundle dissection method, with TractSeg consistently resulting in high reproducibility for all sources of variation. Within a method, most pathways show similar patterns of reproducibility. For example, for TractSeg and Xtract all pathways indicate high RESCAN, DIR(1 and 2) and BVAL reproducibility, but are most sensitive to RES, with RES2 showing more variation than RES1. Additionally, Dice overlap shows some variation across pathways, for example CST and UF generally have higher overlap than OR, IFO, and AF, although trends are different for different workflows.

**Figure 3.**
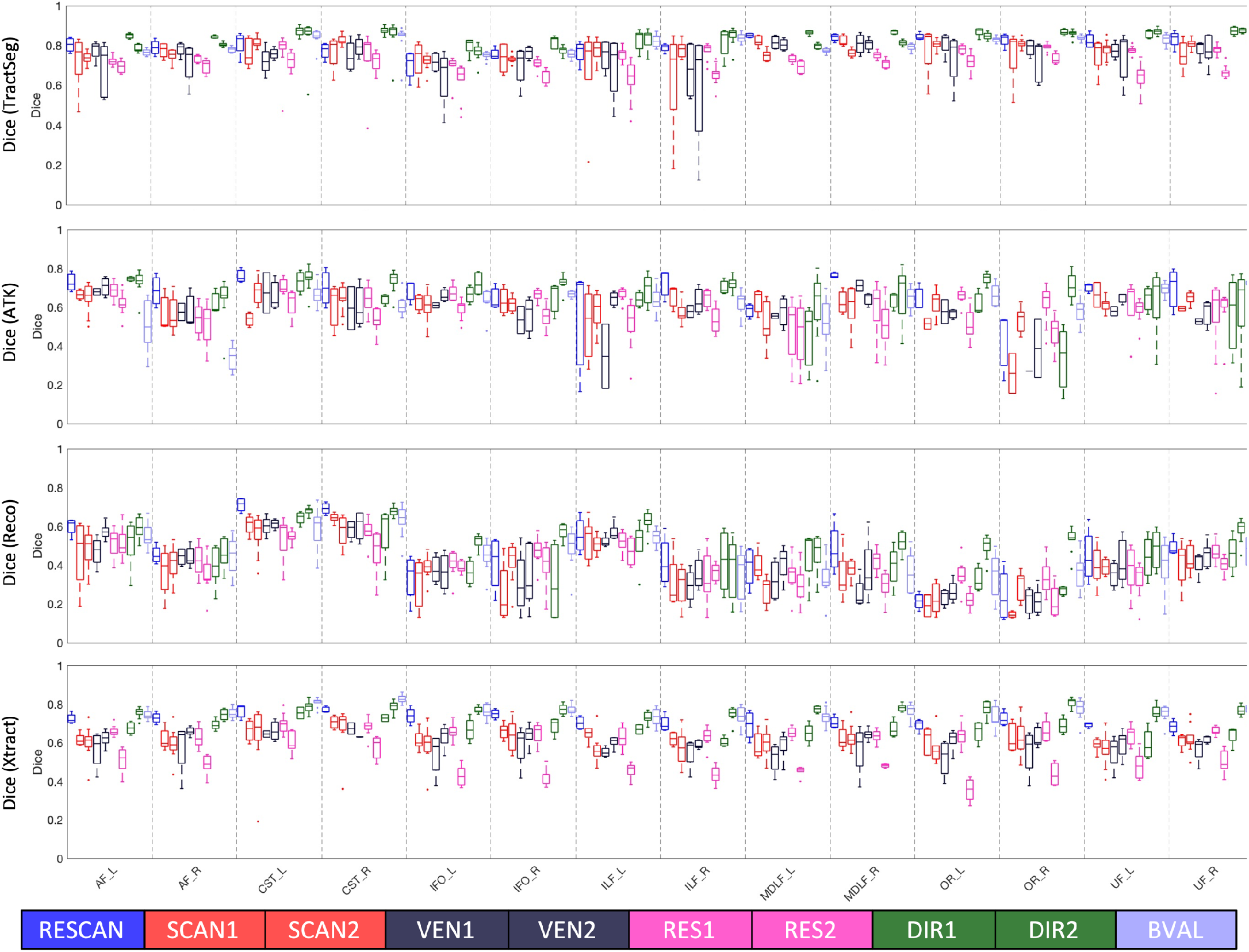
Reproducibility is dependent on all investigated effects, and varies by pathway and by dissection method. Effects of scan-rescan (RESCAN; blue), scanners (SCAN1, SCAN2; red), vendor (VEN1, VEN2; dark purple), resolution (RES1, RES2; pink), diffusion directions (DIR1, DIR2; green) and b-value (BVAL; light purple) on dice overlap coefficient for individual bundles. Results are shown for 14 fiber bundles that are common to each tractography workflow. Please see Appendix for bundle abbreviations.

The results of the Dice overlap coefficient-analysis for each method is shown in **Figure 4**, but condensed across all pathways within a given bundle segmentation method. Similar trends are observed as in **Figure 3**, with TractSeg consistently indicating the highest Dice overlap, and all methods indicating moderate-to-good overall overlap for most pathways. In general, the largest differences are observed when changing resolution, with changes due to RES2 resulting in larger differences than RES1. Following this, differences across vendors (VEN1 more different than VEN2 comparisons) are greater than across scanners (for both SCAN1 and SCAN2), which are greater than the inherently stochastic nature of RESCAN variability. Finally, differences caused by DIR (1 and 2) and BVAL are on the level of, or even less than, those caused by RESCAN, with the notable exception of ATK, which utilizes a reconstruction method and tractography propagation inherently dependent on diffusion sensitization.

**Figure 4.**
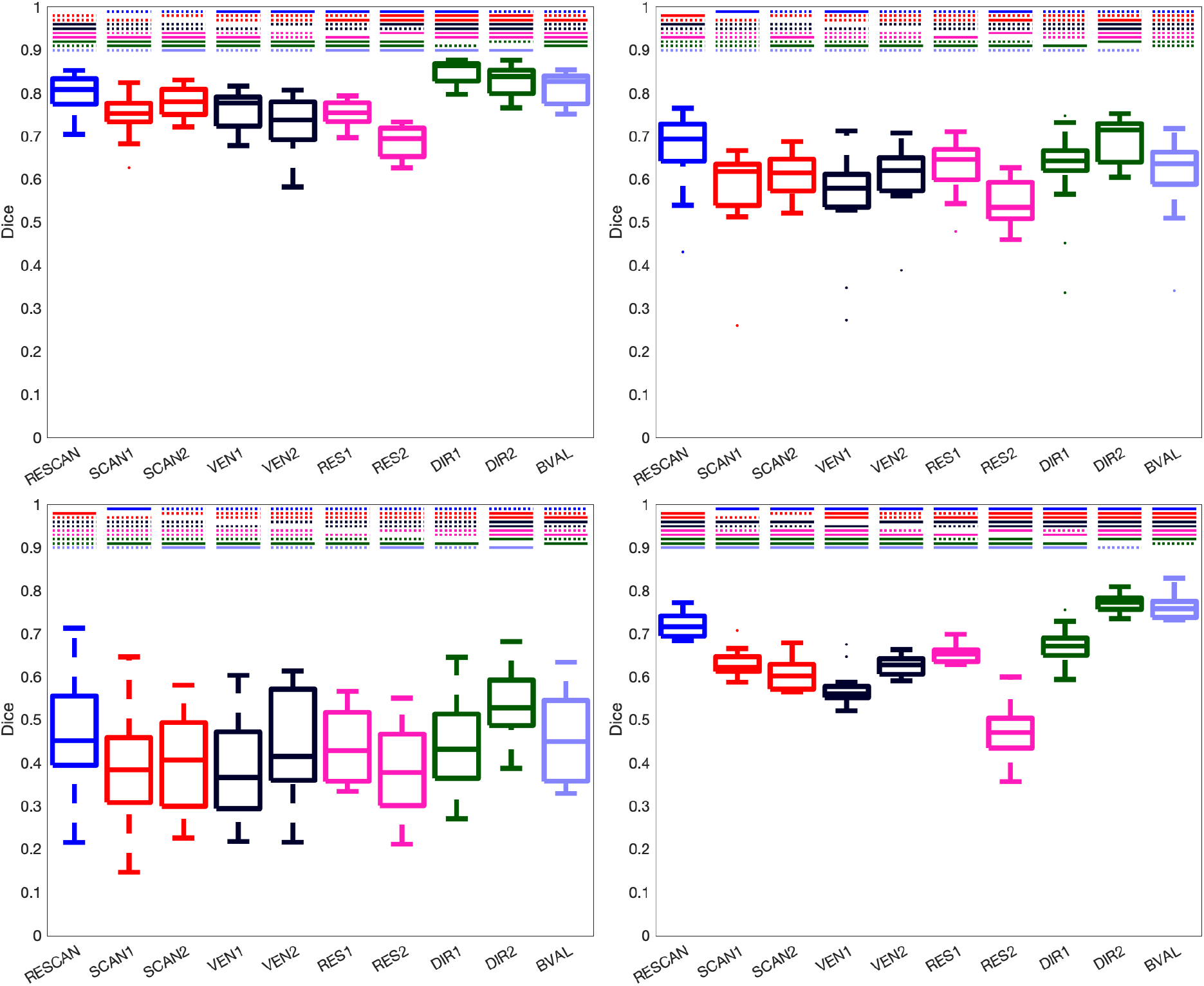
Reproducibility is dependent upon all investigated effects, and each bundle segmentation methods is affected differently. Effects of scan-rescan (RESCAN; blue), scanners (SCAN1, SCAN2; red), vendor (VEN1, VEN2; dark purple), resolution (RES1, RES2; pink), diffusion directions (DIR1, DIR2; green) and b-value (BVAL; light purple) on dice overlap coefficient for all fiber bundles dissected using each technique. For each, a Wilcoxon signed rank test is performed to investigate differences in effects. Statistically significant results (p<.05/45/4 comparisons) are shown as a solid line, and those not reaching statistical significance are shown as dashed line. Tractseg (top-left), ATK (top-right), Reco (bottom-left), and Xtract (bottom-right).

### Localization of Variation

**Figure 5** visualizes locations of tractography bundle segmentation agreement (or consistency), and where it disagrees (variability) as hot and cold colormaps, respectively. Agreement and disagreement are averaged across all subjects and shown for all sources of variation. For display, we have chosen an example pathway that is highly reproducible (the AF from TractSeg) and one which displayed lower reproducibility (the SLFII from Xtract). For the highly reproducible pathway, all sources of variation show very similar results. The agreement is very high throughout the entire pathway (hot colors), and percent-disagreement remains fairly low (black and dark blue colors). This means that when two bundles disagree, the disagreement is largely randomly distributed, rather than a *consistent* localized bias introduced by a certain source of variation – an effect which would show up as a consistent disagreement (i.e. a high percent-disagreement). Disagreement tends to occur at the periphery, or boundaries, of the pathway, in particular at the gray-white matter junction, and within individual gyri.

**Figure 5.**
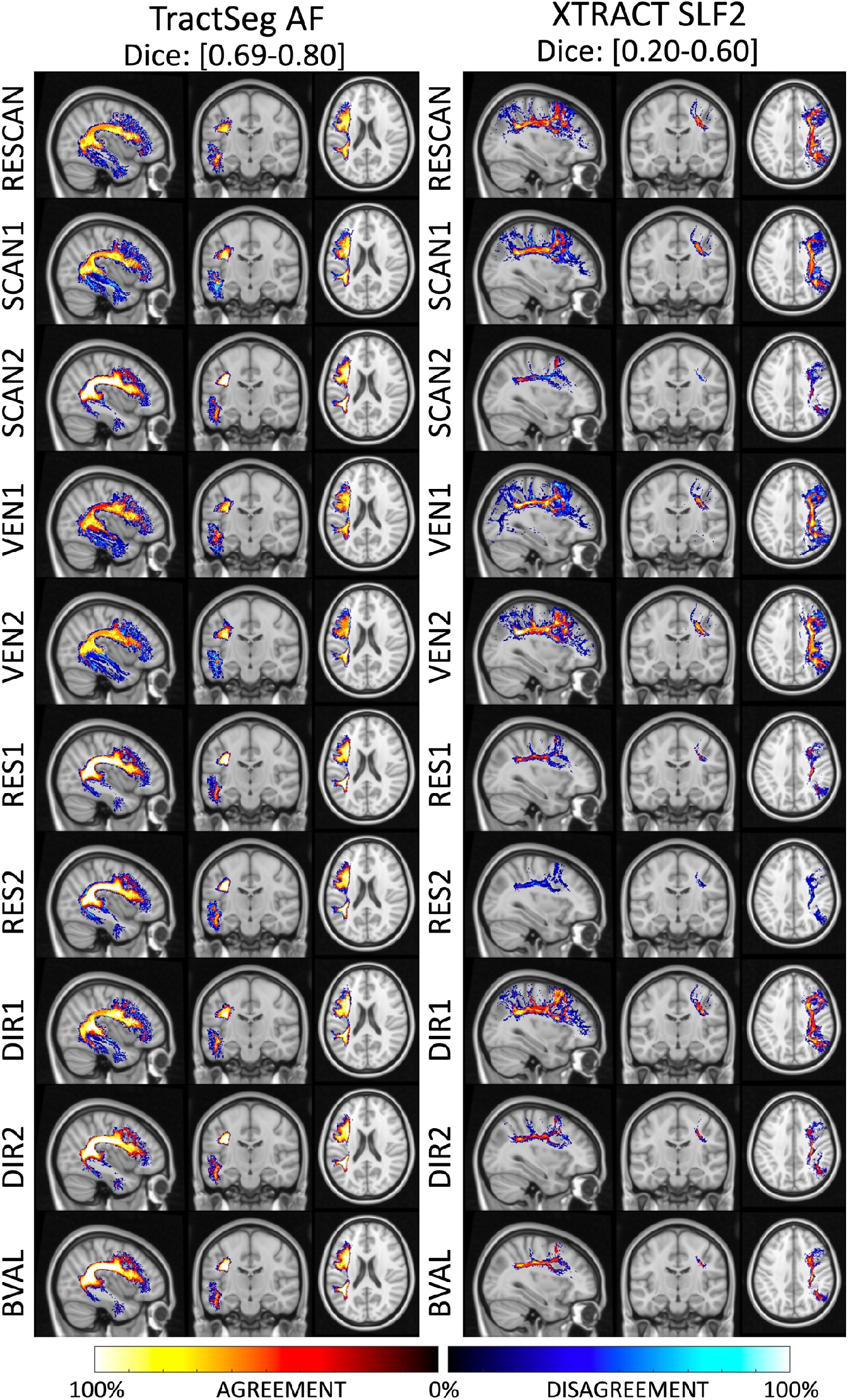
Locations of agreement and disagreement across effects. Maps are computed by overlaying (for each source of variation), maps of where there is overlap (i.e. agreement) and non-overlap (disagreement), averaged across all subjects. For each effect, the percent agreement indicates areas where a pathway is consistently located and is shown using a “hot” colormap, while the percent disagreement indicates areas without consistent overlap and is shown using a “cold” colormap. Results are shown for a highly reproducible pathway (AF_L dissected using TractSeg) and for a less reproducible pathway (SLF2 dissected using XTRACT). Note that even though disagreement is abundant, it does not consistently occur (i.e., % disagreement remains low; black and dark blue) suggesting no systematic bias due to effects, and disagreements are largely attributed to the stochastic nature of the tractography and dissection process.

For the less reproducible pathway, the agreement is moderate to high in the dense core, or center, of the pathway in the deep white matter. Again, disagreements are at the edges, and prominent at the white matter and gray matter boundary. However, even though disagreement is more noticeable, the percent-disagreement remains low, indicating random disagreement as opposed to a consistent bias in the spatial location of this pathway. In this case, sources of variation from SCAN2 and RES2 and VEN1 are more noticeable as a larger source of variation, in agreement with quantitative results.

### Variation of shape features

**Figure 6** shows the RESCAN reproducibility of shape features as measured by MAPE, for all features and all pathways, visualized in decreasing reproducibility. In agreement with Dice, TractSeg has higher overall reproducibility, with most features and most pathways below 10% MAPE. Similarly, ATK and Reco are able to reproducibly characterize most features of most pathways with high consistency. In general, reproducibility of features follows similar order across all methods, with features of Curl, Length, Span, and Diameter highly reproducible, and those of surface area, volume, and end area less so. Additionally, reproducibility is highly dependent on pathway, with clear variation depending upon the bundle being analyzed.

**Figure 6.**
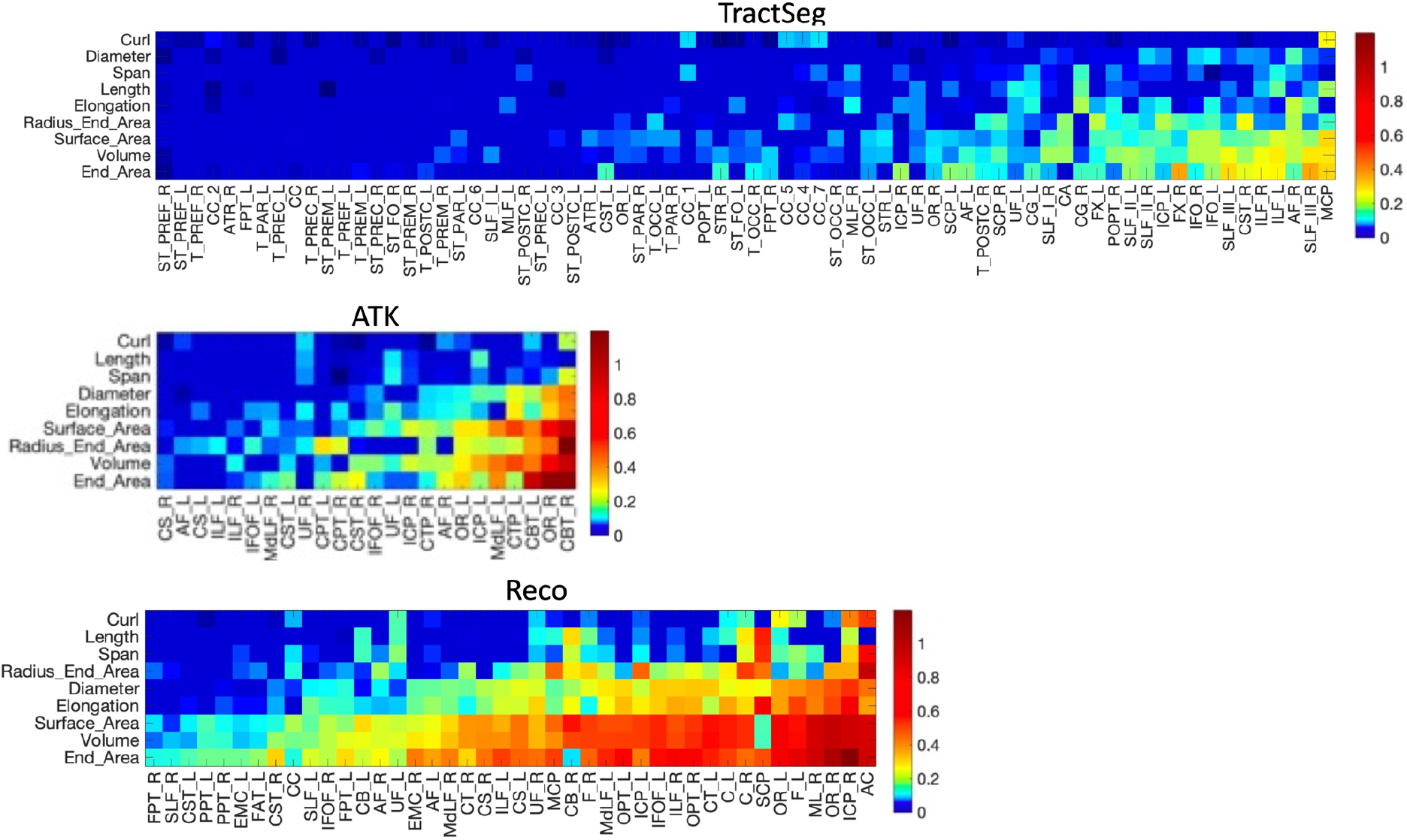
Reproducibility of pathway shape features depends on pathway and bundle dissection method. Reproducibility is shown as a MAPE for each tractography segmentation method. For each method, the features are ordered (from top to bottom) from lowest to highest average MAPE, and pathways are similarly ordered (from left to right) from lowest to highest average MAPE. Note that the colormap is nonlinear to better highlight MAPE between 0-0.10. Many shape features are highly reproducible, and with differences across pathways and bundle dissection methods. Please see Appendix for bundle abbreviations.

**Figure 7** summarizes the MAPE of different features across different sources of variation. Again, Curl, Length, and Span are highly reproducible across all effects, with MAPE always below 10%, and surface area and volume result in higher MAPE. Trends are the same as those observed for Dice overlap, with generally larger differences due to resolution and vendor acquisition effects (RES 1 and 2, VEN 1 and 2), followed by scanner effects (SCAN1 showing the largest variation).

**Figure 7.**
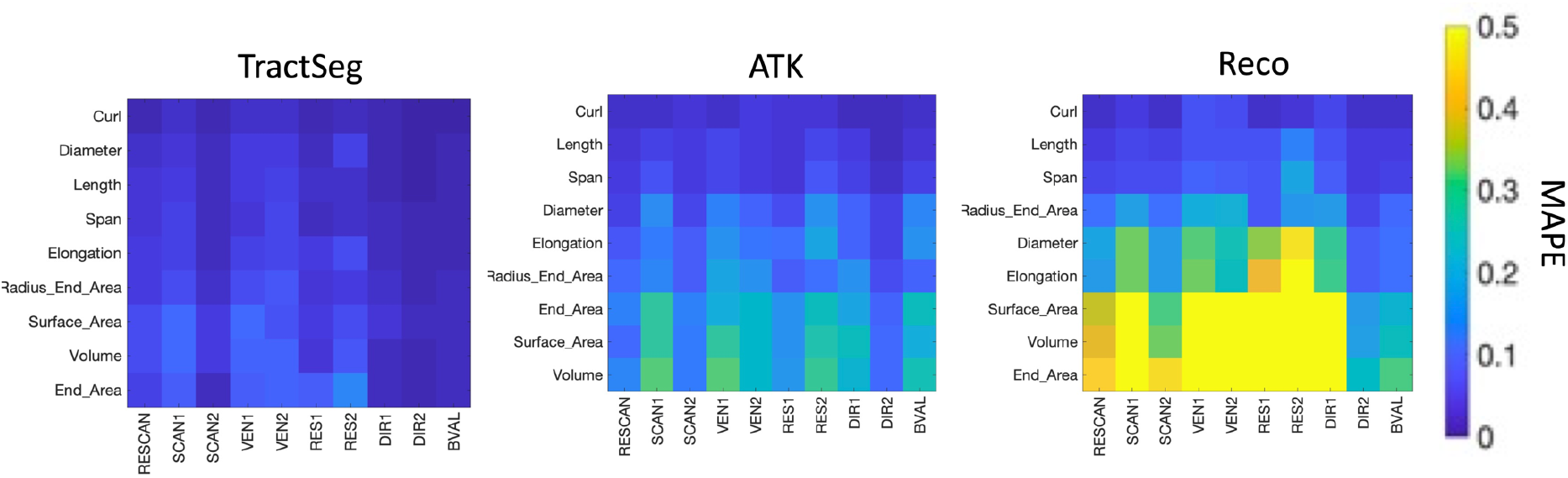
Variability of shape features is influenced by scanner, vendor, acquisition, and b-value. Variability is shown as MAPE for each TractSeg, ATK, and Reco methods, for scan-rescan (RESCAN), scanners (SCAN1, SCAN2), vendor (VEN1, VEN2), resolution (RES1, RES2), diffusion directions (DIR1, DIR2) and b-value (BVAL). Values shown are averaged across all pathways within a bundle dissection method. Shape features are ordered (from top to bottom) from lowest to highest average MAPE. Many shape features are highly reproducible, and MAPE is influenced by all effects investigated.

To look for systematic differences introduced in the quantification of features, we calculate the mean percent variation (i.e., the signed value of MAPE), across all sources of variation, for all features (across all bundles). **Figure 8** shows that most effects do not significantly bias bundle shape measures. For example, nearly all features derived from TractSeg are within a 10% variation and largely centered on 0. However, RES2 and VEN2 do introduce a small, but consistent, bias, in measures of surface area, end area, and volume (in this case, the higher resolution results in smaller values). Similarly, for ATK, a bias is observed in the opposite direction for the same features for effects of acquisition resolution. Additionally, b-value introduces a significant bias for ATK, with the higher b-value scan resulting in larger quantitative values for these features. Reco, in agreement with previous figures, has a much wider range of variation, and larger effects due to acquisition for features of Diameter, Surface Area, End Areas and Volume. Thus, different sources of variation may bias quantitative extraction of shape features, and bias them differently for different bundle segmentation methods.

**Figure 8.**
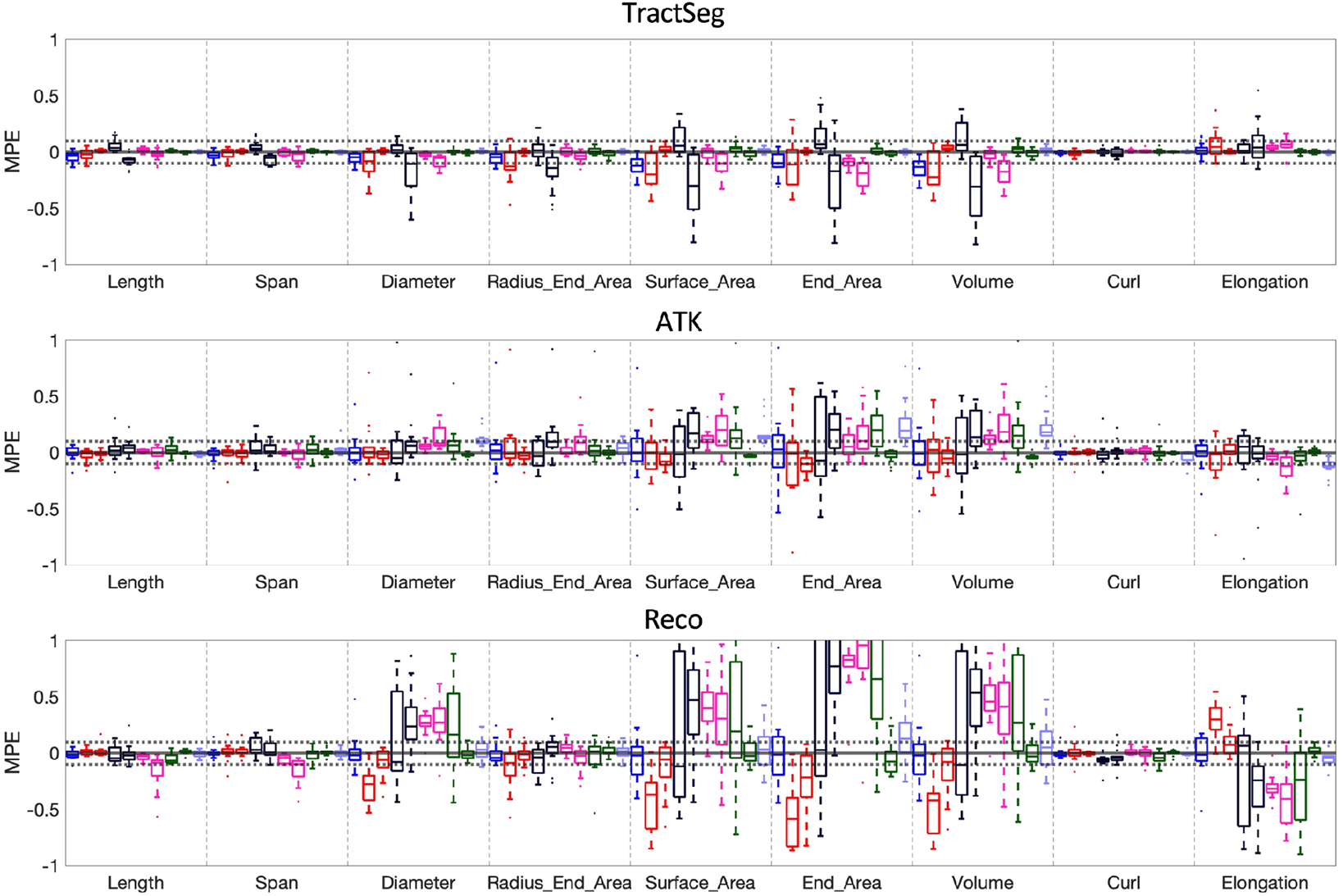
Sources of variation may introduce bias in shape features. The mean percent variation (MPV), i.e., the signed MAPE, is shown for each bundle segmentation method, for all features, with the distribution across fiber pathways. A distribution not centered on 0 suggests systematic differences introduced by the given effect. For interpretation, RESCAN (repeat 2 – repeat 1), SCAN1 (Philips Achieva scanner 2 – Philips Achieva scanner 1), SCAN2 (Siemens Connectome standard acquisition – Siemens Prisma standard acquisition, VEN1 (GE Discovery - Philips Achieva), VEN2 (Siemens Skyra - Philips Achieva), RES1 (Prisma state-of-the-art 30 directions - Prisma standard acquisition), RES2 (Connectom state-of-the-art 30 directions - Connectom standard acquisition), DIR1 (Philips Achieva 96 directions – Philips Achieva 32 directions), DIR2 (Prisma state-of-the-art 60 directions - Prisma state-of-the-art 30 directions), BVAL (Prisma standard-acquisition b=3000 - Prisma standard-acquisition b=1000)

### Variation across bundle segmentation methods

Next, we compared the agreement of the same bundle, but across different bundle segmentation methods. **Figure 9** shows the Dice overlap for 14 common bundles, comparing each method to every other. There is a low-to-moderate agreement, with Dice overlap values between 0.1-0.5 for all pathways. In general, ATK was most similar to TractSeg and Reco for most bundles (with some exceptions), while Xtract was most dissimilar to all others. The AF, ILF, and MDLF, were the most dissimilar across methods.

**Figure 9.**
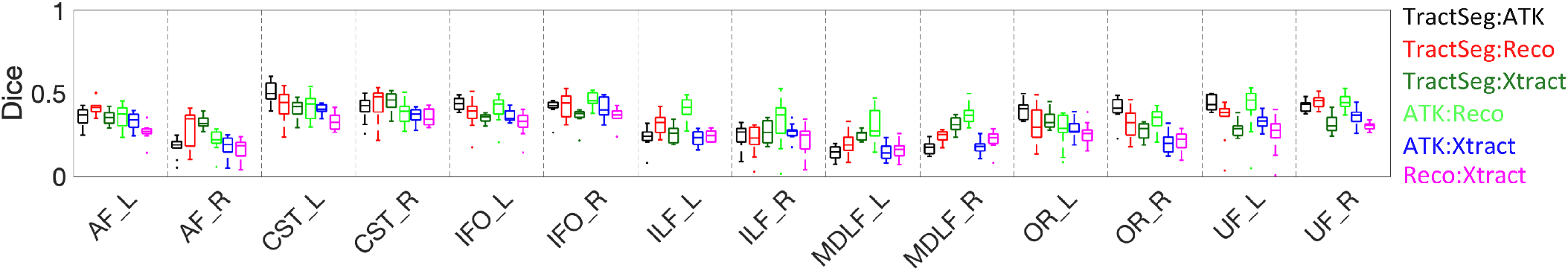
Different workflows result in low-to-moderate Dice overlap of the when measuring agreement between different bundle dissection methods. Please

**Figure 10** visualizes where agreement and disagreement occurs across bundle segmentation methods, with example-pathways AF and OR. Here, while most of the core agrees across methods, there is also a *consistent* disagreement across methods, particularly in the thickness of the bundle and in the regions of the temporal lobe for the AF and connections in the occipital lobe for the OR. Thus, instead of random differences due to noise, differences across methods are reproducible disagreement, likely caused by fundamental differences in the segmentation technique and structure to be segmented.

**Figure 10.**
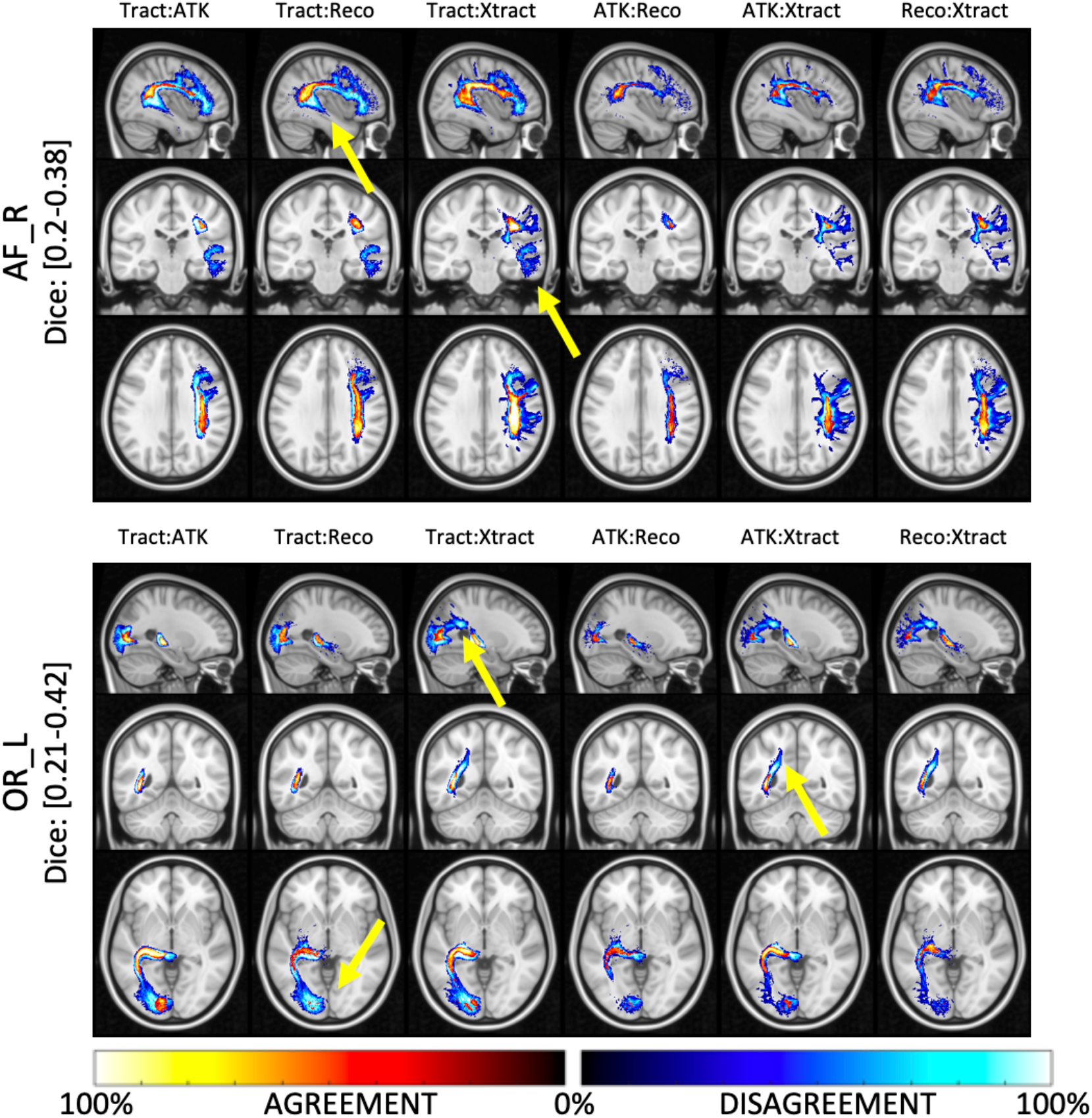
Locations of agreement and disagreement across bundle dissection methods. For each comparison, percent agreement indicates areas where methods agree in space and is shown using a “hot” colormap, while percent disagreement indicates areas where disagreement occurs and is shown using a “cold” colormap. Results are shown for two example pathways (AF_R and OR_L). Here, there are areas of high % disagreement between methods, indicating a consistent and reproducible difference between bundle dissection methods (highlighted by yellow arrows).

### Variation in diffusion MRI microstructure measures

We next investigate reproducibility of microstructure measures due to the aforementioned sources of variation, and tractography variation. **Figure 11** shows the MAPE of FA for all four bundle segmentation methods. In all cases, the standard-color boxplots are variations due to the queried source of variation alone, whereas the darker-shaded boxplots are due to the source of variation *and* the added variation of tractography variation. Most notably, the MAPE due to RESCAN, SCAN, VEN, DIR, and BVAL alone are highly similar for all segmentation methods, with only minor differences due to the slightly different representations of the pathways (**Figure 9**). These results are in line with the literature, with variation <3% for SCAN rescan [24, 25, 58], with 5-15% due to scanner and vendor effects [40, 41], and as much as 10% due to differences in acquisition and diffusion sensitization [33, 40, 74]. Notably, the added variation due to tractography does indeed increase differences in FA (as indicated by a solid horizontal line) in many cases, although the % increase in variation is on average <5%.

**Figure 11.**
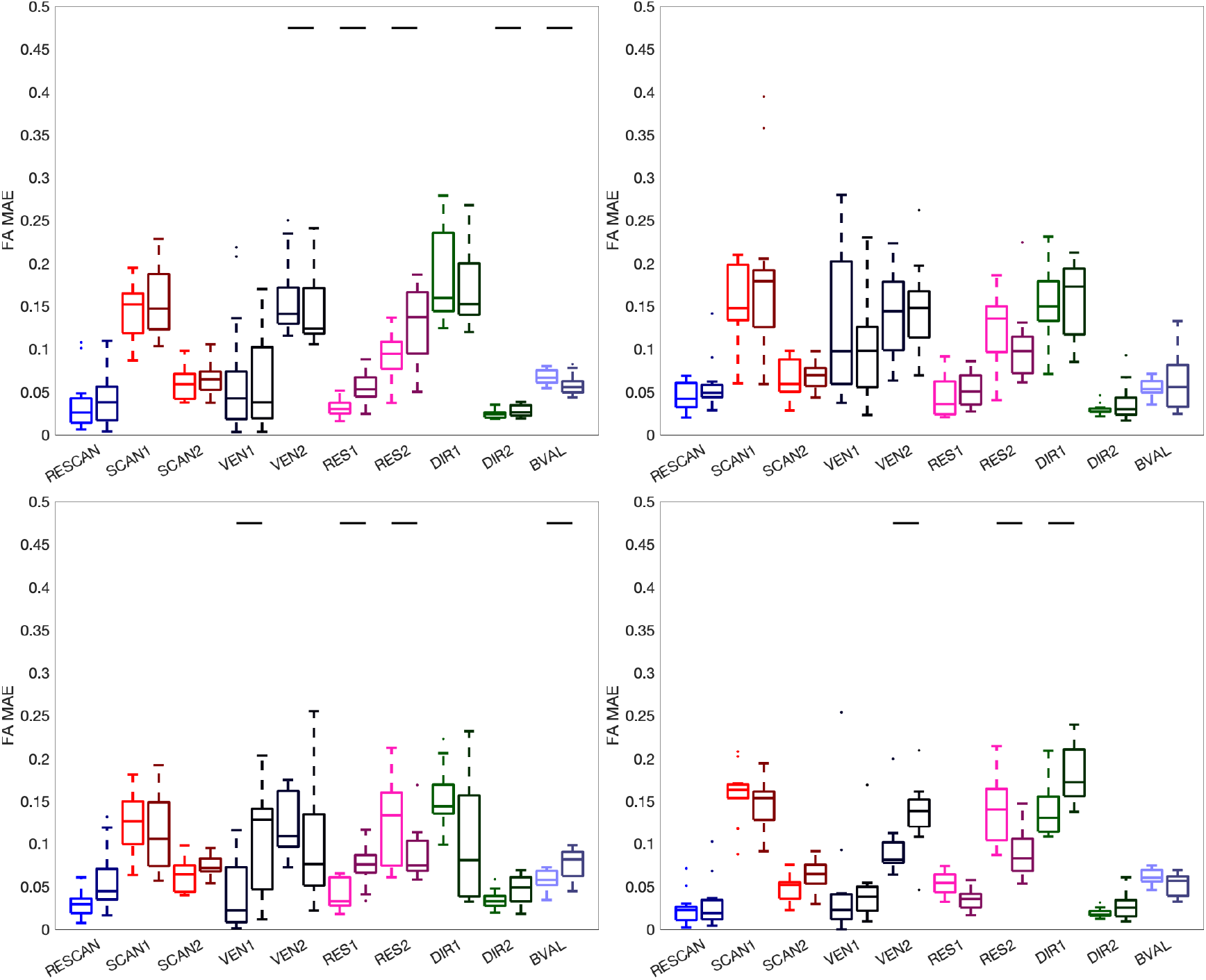
Variation of FA. Effects of scan-rescan (RESCAN; blue), scanners (SCAN1, SCAN2; red), vendor (VEN1, VEN2; dark purple), resolution (RES1, RES2; pink), diffusion directions (DIR1, DIR2; green) and b-value (BVAL; light purple) on MAPE of the FA for all fiber bundles dissected using each technique. The left boxplots are indictive of the variability inherent due to each effect, whereas the darker-hued (right) boxplots indicate the added variability due to differences in tractograms. For each, a Wilcoxon signed rank test is performed to investigate whether tractography adds to (or removes) significant variance to this metric, and statistical significance is indicated by a solid black line.

**Figure 12** shows the MAPE of MD for different sources of variation. Most noticeable, MD is highly different when calculated using two different b-values, as expected [3, 25, 32, 75, 76], followed by differences due to vendors. Differences across RESCAN, SCAN, RES, and DIR are typically <5%. Again, the use of tractography adds to this variance, although on 3% or less on average.

**Figure 12.**
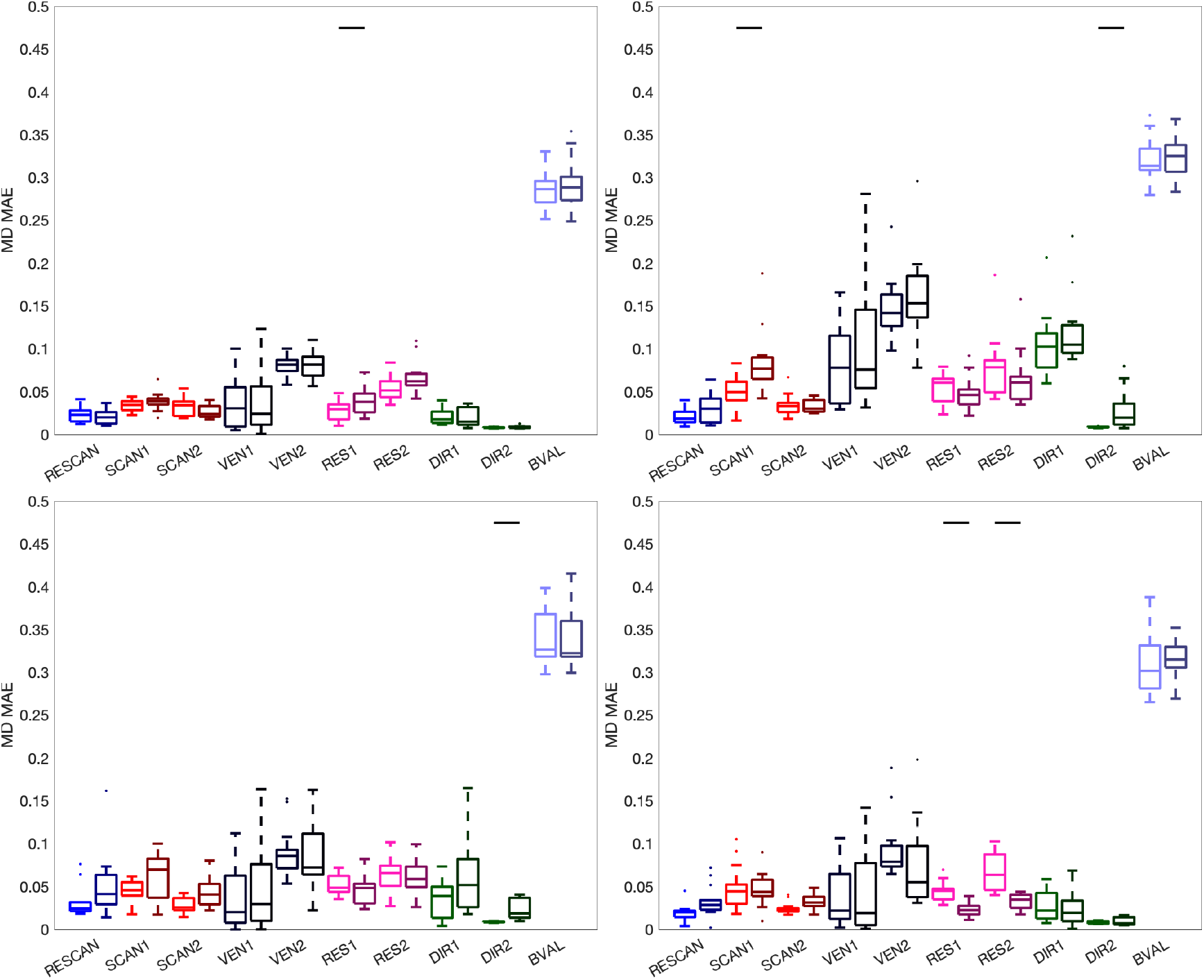
Variation of MD. Effects of scan-rescan (RESCAN; blue), scanners (SCAN1, SCAN2; red), vendor (VEN1, VEN2; dark purple), resolution (RES1, RES2; pink), diffusion directions (DIR1, DIR2; green) and b-value (BVAL; light purple) on MAPE of the MD for all fiber bundles dissected using each technique. The left boxplots are indictive of the variability inherent due to each effect, whereas the darker-hued (right) boxplots indicate the added variability due to differences in tractograms. For each, a Wilcoxon signed rank test is performed to investigate whether tractography adds to (or removes) significant variance to this metric, and statistical significance is indicated by a solid black line.

## Discussion

The primary focus of this work was to study variability of diffusion fiber tractography bundle segmentation, performing the same analysis on different datasets on different scanners or with different acquisition protocols. For the databases investigated here, we have shown that the process of tractography bundle segmentation shows significant variation across different acquisition resolution and across different vendors, with less, albeit significant, variation across scanners and across diffusion sensitization. Variation is indeed expected when scanning the same subject twice, with all other experimental parameters constant, due to imaging noise and the stochastic nature of the tractography process, however, these additional sources of variation add potential confounds to tractography analysis that may bias measurements, limit aggregation of datasets, and hinder direct interpretation and meta-analysis of different results across studies. While the primary focus was on variation due to vendor and scanner effects, acquisition effects, and b-value effects, we also show the most bundle segmentation workflows are highly reproducible when running the same analysis on data acquired in different sessions, but with the same scanner and protocol.

It is well-known that microstructural features at different sites and with different protocols are not immediately comparable, and in fact significantly biased due to various effects. However, the process of tractography is largely dependent upon fiber orientation estimates, rather than features of the signal magnitude directly (i.e., MD/FA), and it is not immediately intuitive that differences in scanners, acquisitions, and b-values may lead to significantly different results. The results of this work suggest that, indeed, the results of tractography and across sites adds variability that must be considered in the interpretation of both microstructural and shape features of these pathways.

### Do we need to harmonize tractography?

“Harmonization” can be considered any effort at reducing variability in quantitative metrics between different databases, scanners, and studies. We have known that the voxel-wise signal varies across sites, scanners, and acquisitions (as evidenced by the multitude of efforts in the literature to study effects on DTI-indices [1-15]) and now confirm that the tractography process itself does as well, and have quantified the extent that tractography contributes to variability. The question becomes “do we need to harmonize tractography?”. The short answer is “yes”, the long answer is: harmonizing likely entails both harmonizing the signal (e.g., FA, MD, RISH measures), harmonizing orientation, reducing effects of resolution, and combining the strengths of different bundle segmentation approaches.

The field of diffusion MRI harmonization has grown in recent years, with significant efforts to make diffusion microstructural measures comparable across sites and scanners [19, 40, 42-44, 47, 48]. Yet, these endeavors have traditionally not considered variability of tractography, which is ultimately influenced at both the local scale of individual voxels and voxel-wise reconstruction as well as a global scale of connecting discrete orientation estimates across the brain.

It is unclear what “harmonizing” tractography may entail. Clearly, consistent orientation estimates are key, but also streamline generation algorithms robust to voxel-sizes, and also segmentation algorithms that are consistently able to identify streamlines belonging to a pathway-of-interest. With the vast array of options to reconstruct orientation, generate streamlines, and segment bundles, it may be impossible to harmonize data in a way that is appropriate for all methods. Some effort has been performed to harmonize fiber orientation estimation specifically across time or across scanners [45, 77-79]. It may be possible that harmonizing the microstructural measures themselves may remove some possible confounds (i.e., if FA is used as a stopping criteria). Similarly, it is possible that the application and process of tractography in a standard space (as performed for XTRACT), or at a standard resolution may remove confounds associated with image resolution. Alternatively, various multi-site methods used for scalar microstructure features, instead of harmonizing bundles of streamlines directly, may be utilized to harmonize features extracted from bundles. Finally, even while there is significant variation, large agreement occurs in the core of reconstructed white matter pathways, and weighting all derived measures and features by tract density, or isolating the trunk of the bundle [7], may remove sources of variation.

Reassuringly, the automated methods considered are fairly robust to these studied sources of variation. Visually, the pathways look remarkably similar across scanners, acquisition, and protocols (**Figure 2**), for all methods. Quantitatively, methods such as TractSeg, which utilize orientation estimates alone, in combination with machine learning techniques in order to map out tract orientation maps, endpoints, and binary segmentations are highly reproducible. Similarly, the other methods, while quantitatively having moderately larger variation, show similar shapes, locations, and connectivity across all effects. A final possible harmonization approach may be to combine the strengths of the various algorithms, rethinking the process of bundle segmentation to possibly utilize some combination of machine learning (TractSeg), and a volume-based extraction prior to streamline generation, followed by atlas-based (ATK, Xtract), or shape-based filtering (Reco) in order to delineate bundles consistently across potential confounds.

### Which confounds impact tractography the most?

It is important to emphasize that we are purposefully not attempting to “rank” algorithms, or suggest that ones are better than others. Even the methods with apparent lower reproducibility of features and shapes are still moderately robust, and different implementations of these algorithms may have yielded different quantitative values. For example, different thresholding could have been applied to both density-based (Xtract) or streamline-based (all others) methods to increase specificity (or vice-versa, specificity), or different whole-brain tractography could have been applied prior to bundle dissection using Recobundles. However, regardless of implementation and choices of hyperparameters, we expect methods to show similar dependencies to the investigated sources of variation.

To our knowledge, this is the first time that multiple sources of variation of tractography have been investigated together. Reproducibility across raters, across algorithms, and across scanners have previously been investigated. Our results allow comparison of the relative impact of changes across sites or scanners, and suggest that, in general spatial resolution leads to the most dramatic differences in resulting tractograms. Less tissue-based partial volume effects within the white matter may facilitate delineation of white matter bundles [80]. Additionally, when quantifying volume overlap and shape features, voxel-wise partial volume effects may cause a higher (or lower) estimate due to the representation of the bundle as a binary volume at the given spatial resolution. Finally, orientation-based partial volume effects are observed with different spatial resolution [30, 81], leading to differences in accuracy of fiber orientation distributions, as well as fundamental differences in common diffusion measures such as FA (which are often used in the tracking process).

The second biggest contributor to variability was vendor differences. Differences across scanners are known to introduce variability due to factors of maximum gradient strength (and hence echo times and repetition times), field strengths, gradient nonlinearities, receive coil sensitivities, software version, and system calibration [19]. Here, we show that differences in vendors are typically greater than that due to different scanners (yet same vendor) alone. Over and above scanner differences, vendors themselves may variations in algorithm choices, algorithms for acquisition, reconstruction, background noise reduction, multi-coil fusion [82, 83], and pulse sequence implementation. Here, we have shown that in addition to inconsistencies in DTI measures across vendors consistently shown in previous studies [84-86] there is also a large inconsistency in tractography volumes and locations due to differences in vendors.

Reassuringly, variation of b-value and number of diffusion directions led to relatively consistent tractography. While it is well-known that angular resolution affects the ability to reconstruct fiber orientations [30, 87-91], most reconstruction methods are robust with as few as 30 directions (or less). Similarly, while reconstruction algorithms are dependent on diffusion sensitization [87, 92], the b-value did not significantly affect tractography results (although does affect quantitative metrics association with DTI).

It is also interesting that the relative magnitude of sources of variation depend on the bundle dissection method. While variability generally decreases from RESCAN, DIR, BVAL, SCAN, then VEN and RES, several notable exceptions occur. ATK is highly sensitive to the b-value. This is likely due to the fact that this automated tractography is reconstructed using Generalized Q-ball Imaging [69], and tracking thresholds are determined by the normalized quantitative anisotropy, which is known to be highly dependent on b-value [93]. In contrast, XTRACT is a probabilistic method based largely on fiber orientation (and its dispersion) alone (from the ball-and-stick model [94]), and different b-values give highly similar results of orientation (although dispersion will vary). XTRACT is also most sensitive to drastic change in resolution, likely caused by the probabilistic nature of the tractography process and subsequent thresholding for segmentation.

### Shape variation and location of variation

This is to the best of our knowledge also the first time that reproducibility of different shape features of tractography has been investigated. While the variation across and within subjects has previously been studied [68], it is important to understand cross-protocol and cross-scanner effects if these features are to be potential biomarkers in health and disease. These shape measures show similar patterns of variability, largest across resolution, vendors, and scanners scanners, and smallest variation across repeats, directions, and b-values. More than variation, different resolutions and b-values can significantly bias measures, for example consistently overestimating volume and surface areas at lower resolutions where more partial volume effects are expected. Depending on tractography method, many features are remarkably robust, with MAPE below 5%, in line with that of microstructure features.

We also investigated locations of differences and similarities by visualizing where there was consistent agreement and disagreement. Importantly, even with differences in acquisition and scanners, methods are able to consistently reproduce the major shape and location of the intended pathway, with differences most frequently occurring at the periphery, or edges, of the pathway, and along the white matter and gray matter interface. While features of shape and geometry may be biased due to sources of variation, these differences do not consistently occur at any one location or place along the pathway.

### Different workflows

Over and above the typically studied sources of variation, we found that differences due to the choice of bundle segmentation workflows are most pronounced. For any given pathway, overlap from one workflow to another was low-to-moderate. This is in part due to the inherent sensitivity/specificity of different algorithms – for example Recobundles will look for clusters exhibiting a certain shape, while Tractseg is based on deep-learned segmentation, and Xtract will be highly dependent on the chosen threshold – but more importantly due to fundamental differences in how the pathway is dissected or defined [55, 95]. For example, the definition of a pathway by one method may be entirely different from another method, including choices in the presence or absence of connections to entire lobes or lobules, or differences in estimated spatial extent of pathways. While differences across methods were larger, they were importantly *consistently* different, meaning that comparing findings using different methods may result in differing conclusions on connectivity or microstructure. Differences between bundle segmentation workflows are also confounded by differences in the entire process of tractography, including differences in modeling, generation of streamlines (i.e., tractography), and bundle segmentation or filtering. Thus, it is intuitive that major differences exist when implementing different standard workflows to study the brain.

### Microstructure variation

Finally, we looked at how much the variation in tractography contribute to the already existing cross-protocol and cross-scanner variation in dMRI measures. For FA, difference across scanners are known to be as much as 5-15% [40, 41], and differences are expected due to different b-values, while scan-rescan reproducibility is high (<5%). The variation in tractography segmentations does indeed statistically significantly increase this variation for most effects, although the increase is typically very small and <5%. Similar results are observed for MD, although most changes are most pronounced for MD across different scanners. Thus, while tractography has the benefit of added specificity over simply propagating atlas-derived regions to subject-space, it does potentially increase variability in these measurements. Although methods such as tract-based spatial statistics [96] have been developed to mitigate these effects, we lose the added benefit of characterizing an index of interest along or within the full trajectory of the pathway.

### Future studies and Limitations

Future studies should investigate additional sources of variation. Manual dissection of fiber bundles gives the dissector the ability to interactively manipulate pathways to their liking [52], and it remains to be seen how this is influenced by scanner and site given the flexibility of this approach. Further, it is unknown whether these variabilities will matter in a clinical setting [97-100], although with the importance of determining pathway boundaries, we hypothesize that the partial volume effects due to acquisition resolution will possibly influence decision making. It is worth investigating the potentially large array of automated bundle segmentation methods that exist, as some are likely more/less appropriate when comparing or combining datasets with different confounds. Additionally, as alternative segmentation methods, or even whole-brain connectome analysis pipelines, are proposed, the use of open-source multi-site multi-subject datasets [101-103] should be encouraged to investigate the successes and limitations of new approaches. Many algorithms for reconstruction and tractography are now able to utilize multiple diffusion shells, and the change in variability and precision of tractography using these techniques compared to isolated diffusion weightings should be compared, but is outside the scope of this work. As along-fiber quantification [6, 7] has proven valuable in the research setting, it would be worthwhile to perform investigations which parallel the current study in order to ask how and where along the bundle differences occur due to different effects. This has been previously investigated, but is largely limited to scan-rescan analysis [7, 101, 104], while the tract-averaged indices are still commonly utilized in neuroimaging studies.

A major limitation of the current study is the limited sample sizes of both datasets due to challenges associated with scanning the same subjects on different scanners and with different protocols. However, there are few multi-site multi-subject databases, and fewer still with varied protocols on the same subjects, whereas here we are able to remove effects across subjects by analyzing only the same subject with different protocols. It is expected that more datasets will become available as big-data and multi-site collaborations become more important to the neuroimaging community, and traveling subjects become common place in order to harmonize across sites. Exemplar open-sourced datasets include that of [105] with N=3 subjects at 20 sites with Prisma scanners and a multi-shell dataset (allowing analysis of RESCAN, SCAN, BVAL, DIR), the traveling human phantom dataset with N=5 subjects at 8 center (SCAN, VEN, DIR), or consortiums such as Pharmacog [106], ADNI [103], HCP [61], or OASIS [107], all with large sample size and repeat scans, but typically limited to RESCAN analysis only or without matched subjects across scanners/vendors/protocols. Because of this, for simplicity, we have chosen two datasets in this study which allow incorporation of all intended sources of variation without compromising readability. While we have looked at a wider range of variability factors than previous studies, we emphasize that these results are based only on two specific databases, and nalysis should be reproduced on other (and new) databases in future work to show generalizability.

Finally, while the primary focus of our study was on variation due to scanner-effects, acquisition-effects, and b-value-effects, our analysis was limited to studying these effects on only four bundle segmentation workflows. We did not implement all existing automated bundle reconstruction pipelines or workflows [7, 53, 56-58, 64, 108-115], however, our selection captures a variety of techniques used to reconstruction bundles, including differences in the use of atlases or regions-of-interest, those based on shape and/or orientation features, machine learning techniques, and differences in the generation of streamlines – a wide variety of vastly different approaches that we consider a strength of this study. To create a tractable parameter space, we have chosen only these four representatives of the wide variety of possible approaches.

Finally, we did not directly perform harmonization techniques in this study. There are dozens of methods available to do this (see [116, 117]), and understanding and characterizing harmonization results across several algorithms would take away from the main focus of this study – which is characterization and ranking of variability across confounds. Further, harmonization would only affect a subset of results (i.e., those looking at FA/MD) as most harmonization approaches leave orientation untouched.

## Conclusion

When investigating connectivity and microstructure of the white matter pathways of the brain using tractography, it is important to understand potential confounds and sources of variation in the process. Here, we find that tractography bundle segmentation results are influenced by the use of different vendors and scanners, and different acquisition choices of resolution, diffusion directions, and diffusion sensitizations, thus results may not be directly comparable when combining data or results across studies. Additionally, different bundle segmentation protocols have different successes/limitations when dealing with sources of variation, and the use of different protocols for bundle segmentation may result in different representations of the same intended pathway. These confounds need to be considered when designing or developing new tractography or bundle dissection algorithms, and when interpreting or combining data across sites.

## Code

Multi-site, multi-scanner, multi-protocol, and multi-subject databases are available for MASIvar (https://openneuro.org/datasets/ds003416) and for MUSHAC (by request). Tractography pipelines are implemented as described by each software package using default parameters for TractSeg (Release 2.3; https://github.com/MIC-DKFZ/TractSeg), ATK (Lct 17 2020 build; http://dsi-studio.labsolver.org), RECO (Dipy 1.2.0 ; https://dipy.org), and XTRACT (FSL 6.0.3; https://fsl.fmrib.ox.ac.uk/fsl/fslwiki/XTRACT). Shape analysis is available in DSI Studio, as Matlab Code (https://github.com/dmitrishastin/tractography_shapes/).

## Acknowledgements

This work was supported by the National Institutes of Health under award numbers R01EB017230, and T32EB001628, and in part by ViSE/VICTR VR3029 and the National Center for Research Resources, Grant UL1 RR024975-01. CMWT was supported by a Sir Henry Wellcome Fellowship (215944/Z/19/Z) and a Veni grant (17331) from the Dutch Research Council (NWO).

## Appendix

The bundles resulting from each segmentation pipeline are given as a list below, with acronyms used in the text.

### Recobundles

Anterior Commisure (AC); Arcuate Fasciculus left (AF_L); Arcuate Fasciculus left (AF_R); Cerebellum left (CB_L); Cerebellum right (CB_R); Cingulum left (C_L); Cingulum right (C_R); Corpus Callosum (CC); Corticospinal Tract left (CST_L); Corticospinal Tract Right (CST_R); Corticostriatal Pathway left (CS_L); Corticostriatal Pathway right (CS_R); Central Tegmental Tract left (CT_L); Central Tegmental Tract right (CT_R); Extreme Capsule left (EMC_L); Extreme Capsule right (EMC_R); Fornix left (F_L); Fornix right (F_R); Frontal Aslant Tract left (FAT_L); Frontal Aslant Tract right (FAT_R); Fronto-pontine tract left (FPT_L); Fronto-pontine tract right (FPT_R); Inferior Cerebellar Peduncle left (ICP_L); Inferior Cerebellar Peduncle right (ICP_R); Inferior Fronto-occipital Fasciculus left (IFOF_L); Inferior Fronto-occipital Fasciculus right (IFOF_R); Inferior Longitudinal Fasciculus left (ILF_L); Inferior Longitudinal Fasciculus right (ILF_R); Middle Cerebellar Peduncle (MCP); Middle Longitudinal Fasciculus left (MdLF_L); Middle Longitudinal Fasciculus right (MdLF_R); Medial Lemniscus left (ML_L); Medial Lemniscus right (ML_R); Occipito Pontine Tract left (OPT_L); Occipito Pontine Tract right (OPT_R); Optic Radiation left (OR_L); Optic Radiation right (OR_R); Parieto Pontine Tract left (PPT_L); Parieto Pontine Tract right (PPT_R); Superior Cerebellar Peduncle (SCP); Superior longitudinal fasciculus left (SLF_L); Superior longitudinal fasciculus right (SLF_R); Uncinate Fasciculus left (UF_L); Uncinate Fasciculus right (UF_R);

### TractSeg

Arcuate fascicle left (AF_L); Arcuate fascicle right (AF_R); Anterior Thalamic Radiation left (ATR_L); Thalamic Radiation right; (ATR_R); Commissure Anterior (CA); Rostrum (CC_1; Genu (CC_2); Rostral body (Premotor) (CC_3); Anterior midbody (Primary Motor) (CC_4); Posterior midbody (Primary Somatosensory) (CC_5); Isthmus (CC_6); Splenium (CC_7); Corpus Callosum – all (CC); Cingulum left (CG_L); Cingulum right (CG_R); Corticospinal tract left (CST_L); Corticospinal tract right (CST_R); Fronto-pontine tract left (FPT_L); Fronto-pontine tract right (FPT_R); Fornix left (FX_L); Fornix right (FX_R); Inferior cerebellar peduncle left (ICP_L); Inferior cerebellar peduncle right (ICP_R); Inferior occipito-frontal fascicle left (IFO_L); Inferior occipito-frontal fascicle right (IFO_R); Inferior longitudinal fascicle left (ILF_L); Inferior longitudinal fascicle right (ILF_R); Middle cerebellar peduncle (MCP); Middle longitudinal fascicle left (MLF_L); Middle longitudinal fascicle right (MLF_R); Optic radiation left (OR_L); Optic radiation right (OR_R); Parieto-occipital pontine left (POPT_L); Parieto-occipital pontine right (POPT_R); Superior cerebellar peduncle left (SCP_L); Superior cerebellar peduncle right (SCP_R); Superior longitudinal fascicle III left SLF_III_L); Superior longitudinal fascicle III right (SLF_III_R); Superior longitudinal fascicle II left (SLF_II_L); Superior longitudinal fascicle II right (SLF_II_R); Superior longitudinal fascicle I left (SLF_I_L); Superior longitudinal fascicle I right (SLF_I_R); Striato-fronto-orbital left (ST_FO_L); Striato-fronto-orbital right (ST_FO_R); Striato-occipital left (ST_OCC_L); Striato-occipital right (ST_OCC_R); Striato-parietal left (ST_PAR_L); Striato-parietal right (ST_PAR_R); Striato-postcentral left (ST_POSTC_L); Striato-postcentral right (ST_POSTC_R); Striato-precentral left (ST_PREC_L); Striato-precentral right (ST_PREC_R); Striato-prefrontal left (ST_PREF_L); Striato-prefrontal right (ST_PREF_R); Striato-premotor left (ST_PREM_L); Striato-premotor right (ST_PREM_R); Thalamo-occipital left (T_OCC_L); Thalamo-occipital right (T_OCC_R); Thalamo-parietal left (T_PAR_L); Thalamo-parietal right (T_PAR_R); Thalamo-postcentral left (T_POSTC_L); Thalamo-postcentral right (T_POSTC_R); Thalamo-precentral left (T_PREC_L); Thalamo-precentral right (T_PREC_R); Thalamo-prefrontal left (T_PREF_L); Thalamo-prefrontal right (T_PREF_R); Thalamo-premotor left (T_PREM_L); Thalamo-premotor right (T_PREM_R); Uncinate fascicle left (UF_L); Uncinate fascicle right (UF_R).

### Xtract

Anterior Commissure (AC); Arcuate Fascile left (AF_L); Arcuate Fascile right (AF_R); Acoustic Radiation left (AR_L); Acoustic Radiation right (AR_R); Anterior Thalamic Radiation left (ATR_L); Anterior Thalamic Radiation right (ATR_R); Cingulum Bundle Dorsal left (CBD_L); Cingulum Bundle Dorsal right (CBD_R); Cingulum Bundle Parahippocampal left (CBP_L); Cingulum Bundle Parahippocampal right (CBP_R); Cingulum Bundle Temporal left (CBT_L); Cingulum Bundle Temporal right (CBT_R); Corticospinal Tract left (CST_L); Corticospinal Tract right (CST_R); Frontal Aslant left (FA_L); Frontal Aslant right (FA_R); Forceps Major (FMA); Forceps Minor (FMI); Fornix left (FX_L); Fornix right (FX_R); Inferior Fronto-occipital Fasciculus left (IFO_L); Inferior Fronto-occipital Fasciculus right (IFO_R); Inferior Longitudinal Fasciculus left (ILF_L); Inferior Longitudinal Fasciculus right (ILF_R); Middle Cerebellar Peduncle (MCP); Medio-Dorsal Longitudinal Fasciculus left (MDLF_L); Medio-Dorsal Longitudinal Fasciculus right (MDLF_R); Optic Radiation left (OR_L); Optic Radiation right (OR_R); Superior Longitudinal Fasciculus 1 left (SLF1_L); Superior Longitudinal Fasciculus 1 right (SLF1_R); Superior Longitudinal Fasciculus 2 left (SLF2_L); Superior Longitudinal Fasciculus 2 right (SLF2_R); Superior Longitudinal Fasciculus 3 left (SLF3_L); Superior Longitudinal Fasciculus 3 right (SLF3_R); Superior Thalamic Radiation left (STR_L); Superior Thalamic Radiation right (STR_R); Uncinate Fasciculus left (UF_L); Uncinate Fasciculus right (UF_R); Vertical Occipital Fasciculus left (VOF_L); Vertical Occipital Fasciculus right (VOF_R).

### ATK

Arcuate_Fasciculus_L (AF_L); Arcuate Fasciculus R (AF_R); Cortico Spinal Tract L (CST_L); Cortico Spinal Tract R (CST_R); Cortico Striatal Pathway L (CS_L); Cortico Striatal Pathway R (CS_R); Corticobulbar Tract L (CBT_L); Corticobulbar Tract R (CBT_R); Corticopontine Tract L (CPT_L); Corticopontine Tract R (CPT_R); Corticothalamic Pathway L (CTP_L); Corticothalamic Pathway R (CTP_R); Inferior Cerebellar Peduncle L (ICP_L); Inferior Cerebellar Peduncle R (ICP_R); Inferior Fronto Occipital Fasciculus L (IFOF_L); Inferior Fronto Occipital Fasciculus R (IFOF_R); Inferior Longitudinal Fasciculus L (ILF_L); Inferior Longitudinal Fasciculus R (ILF_R); Optic Radiation L (OR_L); Optic Radiation R (OR_R); Middle Longitudinal Fasciculus L (MdLF_L); Middle Longitudinal Fasciculus R (MdLF_R); Uncinate Fasciculus L (UF_L); Uncinate Fasciculus R (UF_R).

## References

1. Alexander, D.C., et al., Imaging brain microstructure with diffusion MRI: practicality and applications. NMR Biomed, 2019. 32(4): p. e3841.

2. Jones, D.K., et al., Microstructural imaging of the human brain with a ‘super-scanner’: 10 key advantages of ultra-strong gradients for diffusion MRI. Neuroimage, 2018. 182: p. 8–38.

3. Novikov, D.S., et al., Quantifying brain microstructure with diffusion MRI: Theory and parameter estimation. NMR Biomed, 2018: p. e3998.

4. Jeurissen, B., et al., Diffusion MRI fiber tractography of the brain. NMR Biomed, 2019. 32(4): p. e3785.

5. Raffelt, D.A., et al., Investigating white matter fibre density and morphology using fixel-based analysis. Neuroimage, 2017. 144(Pt A): p. 58–73.

6. Chamberland, M., et al., Dimensionality reduction of diffusion MRI measures for improved tractometry of the human brain. Neuroimage, 2019. 200: p. 89–100.

7. Yeatman, J.D., et al., Tract profiles of white matter properties: automating fiber-tract quantification. PLoS One, 2012. 7(11): p. e49790.

8. Maffei, C., S. Sarubbo, and J. Jovicich, Diffusion-based tractography atlas of the human acoustic radiation. Sci Rep, 2019. 9(1): p. 4046.

9. Forkel, S.J., et al., The anatomy of fronto-occipital connections from early blunt dissections to contemporary tractography. Cortex, 2014. 56: p. 73–84.

10. Hau, J., et al., Revisiting the human uncinate fasciculus, its subcomponents and asymmetries with stem-based tractography and microdissection validation. Brain Struct Funct, 2017. 222(4): p. 1645–1662.

11. Hau, J., et al., Cortical Terminations of the Inferior Fronto-Occipital and Uncinate Fasciculi: Anatomical Stem-Based Virtual Dissection. Front Neuroanat, 2016. 10: p. 58.

12. Sarubbo, S., et al., Uncovering the inferior fronto-occipital fascicle and its topological organization in non-human primates: the missing connection for language evolution. Brain Struct Funct, 2019. 224(4): p. 1553–1567.

13. Sarubbo, S., et al., Frontal terminations for the inferior fronto-occipital fascicle: anatomical dissection, DTI study and functional considerations on a multi-component bundle. Brain Struct Funct, 2013. 218(1): p. 21–37.

14. Neubert, F.X., et al., Connectivity reveals relationship of brain areas for reward-guided learning and decision making in human and monkey frontal cortex. Proc Natl Acad Sci U S A, 2015. 112(20): p. E2695–704.

15. Neubert, F.X., et al., Comparison of human ventral frontal cortex areas for cognitive control and language with areas in monkey frontal cortex. Neuron, 2014. 81(3): p. 700–13.

16. Mars, R.B., et al., Connectivity-based subdivisions of the human right “temporoparietal junction area”: evidence for different areas participating in different cortical networks. Cereb Cortex, 2012. 22(8): p. 1894–903.

17. Pierpaoli, C., et al., Diffusion Tensor MR Imaging of the Human Brain. Radiology, 1996. 201: p. 637–648.

18. Prohl, A.K., et al., Reproducibility of Structural and Diffusion Tensor Imaging in the TACERN Multi-Center Study. Frontiers in integrative neuroscience, 2019. 13: p. 24–24.

19. Mirzaalian, H., et al., Inter-site and inter-scanner diffusion MRI data harmonization. Neuroimage, 2016. 135: p. 311–23.

20. Magnotta, V.A., et al., Multicenter reliability of diffusion tensor imaging. Brain connectivity, 2012. 2(6): p. 345–355.

21. Landman, B.A., et al., Multi-parametric neuroimaging reproducibility: a 3-T resource study. Neuroimage, 2011. 54(4): p. 2854–66.

22. Teipel, S.J., et al., Multicenter stability of diffusion tensor imaging measures: A European clinical and physical phantom study. Psychiatry Research: Neuroimaging, 2011. 194(3): p. 363–371.

23. Vollmar, C., et al., Identical, but not the same: Intra-site and inter-site reproducibility of fractional anisotropy measures on two 3.0T scanners. NeuroImage, 2010. 51(4): p. 1384–1394.

24. Farrell, J.A., et al., Effects of signal-to-noise ratio on the accuracy and reproducibility of diffusion tensor imaging-derived fractional anisotropy, mean diffusivity, and principal eigenvector measurements at 1.5 T. J Magn Reson Imaging, 2007. 26(3): p. 756–67.

25. Landman, B.A., et al., Effects of diffusion weighting schemes on the reproducibility of DTI-derived fractional anisotropy, mean diffusivity, and principal eigenvector measurements at 1.5T. Neuroimage, 2007. 36(4): p. 1123–38.

26. Heiervang, E., et al., Between session reproducibility and between subject variability of diffusion MR and tractography measures. Neuroimage, 2006. 33(3): p. 867–77.

27. Pfefferbaum, A., E. Adalsteinsson, and E.V. Sullivan, Replicability of diffusion tensor imaging measurements of fractional anisotropy and trace in brain. J Magn Reson Imaging, 2003. 18(4): p. 427–33.

28. Jones, D.K., Determining and visualizing uncertainty in estimates of fiber orientation from diffusion tensor MRI. Magn Reson Med, 2003. 49(1): p. 7–12.

29. Lori, N.F., et al., Diffusion tensor fiber tracking of human brain connectivity: aquisition methods, reliability analysis and biological results. NMR Biomed, 2002. 15(7-8): p. 494–515.

30. Jones, R., et al., Insight into the fundamental trade-offs of diffusion MRI from polarization-sensitive optical coherence tomography in ex vivo human brain. Neuroimage, 2020: p. 116704.

31. Papinutto, N.D., F. Maule, and J. Jovicich, Reproducibility and biases in high field brain diffusion MRI: An evaluation of acquisition and analysis variables. Magn Reson Imaging, 2013. 31(6): p. 827–39.

32. Jones, D.K., The effect of gradient sampling schemes on measures derived from diffusion tensor MRI: a Monte Carlo study. Magn Reson Med, 2004. 51(4): p. 807–15.

33. Jones, D.K. and P.J. Basser, “Squashing peanuts and smashing pumpkins”: how noise distorts diffusion-weighted MR data. Magn Reson Med, 2004. 52(5): p. 979–93.

34. Jones, D.K., et al., What happens when nine different groups analyze the same DT-MRI data set using voxel-based methods. 2007.

35. Chang, L.C., D.K. Jones, and C. Pierpaoli, RESTORE: robust estimation of tensors by outlier rejection. Magn Reson Med, 2005. 53(5): p. 1088–95.

36. Jones, D.K., T.R. Knosche, and R. Turner, White matter integrity, fiber count, and other fallacies: the do’s and don’ts of diffusion MRI. Neuroimage, 2013. 73: p. 239–54.

37. Jones, D.K., Precision and accuracy in diffusion tensor magnetic resonance imaging. Top Magn Reson Imaging, 2010. 21(2): p. 87–99.

38. Jones, D.K. and M. Cercignani, Twenty-five pitfalls in the analysis of diffusion MRI data. NMR Biomed, 2010. 23(7): p. 803–20.

39. Jones, D.K., M.A. Horsfield, and A. Simmons, Optimal Strategies for Measuring Diffusion in Anisotropic Systems by Magnetic Resonance Imaging. Magnetic Resonance in Medicine, 1999. 42: p. 515–525.

40. Ning, L., et al., Cross-scanner and cross-protocol multi-shell diffusion MRI data harmonization: Algorithms and results. NeuroImage, 2020. 221: p. 117128.

41. Tax, C.M., et al., Cross-scanner and cross-protocol diffusion MRI data harmonisation: A benchmark database and evaluation of algorithms. Neuroimage, 2019. 195: p. 285–299.

42. Zhong, J., et al., Inter-site harmonization based on dual generative adversarial networks for diffusion tensor imaging: application to neonatal white matter development. Biomed Eng Online, 2020. 19(1): p. 4.

43. Cetin Karayumak, S., et al., Retrospective harmonization of multi-site diffusion MRI data acquired with different acquisition parameters. Neuroimage, 2019. 184: p. 180–200.

44. Huynh, K.M., et al., Multi-Site Harmonization of Diffusion MRI Data via Method of Moments. IEEE Trans Med Imaging, 2019. 38(7): p. 1599–1609.

45. Vishwesh Nath, K.G.S., Prasanna Parvathaneni, Colin B. Hansen, Allison E. Hainline, Camilo Bermudez, Samuel Remedios, Justin A. Blaber, Vaibhav Janve, Yurui Gao, Iwona Stepniewska, Baxter P. Rogers, Allen T. Newton, Taylor Davis, Jeff Luci, Adam W. Anderson, Bennett A. Landman. Inter-Scanner Harmonization of High Angular Resolution DW-MRI using Null Space Deep Learning. in MICCAI-CDMRI. 2018. Granada, Spain.

46. Yu, M., et al., Statistical harmonization corrects site effects in functional connectivity measurements from multi-site fMRI data. Hum Brain Mapp, 2018. 39(11): p. 4213–4227.

47. Mirzaalian, H., et al., Multi-site harmonization of diffusion MRI data in a registration framework. Brain Imaging Behav, 2018. 12(1): p. 284–295.

48. Fortin, J.P., et al., Harmonization of multi-site diffusion tensor imaging data. Neuroimage, 2017. 161: p. 149–170.

49. Pestilli, F., et al., Evaluation and statistical inference for human connectomes. Nature Methods, 2014. 11(10): p. 1058–1063.

50. Nath, V., et al., Tractography reproducibility challenge with empirical data (TraCED): The 2017 ISMRM diffusion study group challenge. J Magn Reson Imaging, 2019.

51. Chamberland, M., C.M.W. Tax, and D.K. Jones, Meyer’s loop tractography for image-guided surgery depends on imaging protocol and hardware. Neuroimage Clin, 2018. 20: p. 458–465.

52. Rheault, F., et al., Tractostorm: The what, why, and how of tractography dissection reproducibility. Hum Brain Mapp, 2020.

53. Vazquez, A., et al., Automatic group-wise whole-brain short association fiber bundle labeling based on clustering and cortical surface information. Biomed Eng Online, 2020. 19(1): p. 42.

54. Guevara, M., et al., Reproducibility of superficial white matter tracts using diffusion-weighted imaging tractography. Neuroimage, 2017. 147: p. 703–725.

55. Schilling, K.G., et al., Tractography dissection variability: what happens when 42 groups dissect 14 white matter bundles on the same dataset? bioRxiv,2020: p. 2020.10.07.321083.

56. Zhang, F., et al., Test–retest reproducibility of white matter parcellation using diffusion MRI tractography fiber clustering. Human Brain Mapping, 2019. 40(10): p. 3041–3057.

57. Guevara, P., et al., Automatic fiber bundle segmentation in massive tractography datasets using a multi-subject bundle atlas. Neuroimage, 2012. 61(4): p. 1083–99.

58. Wakana, S., et al., Reproducibility of quantitative tractography methods applied to cerebral white matter. Neuroimage, 2007. 36(3): p. 630–44.

59. Cai, L.Y., et al., MASiVar: Multisite, Multiscanner, and Multisubject Acquisitions for Studying Variability in Diffusion Weighted Magnetic Resonance Imaging. bioRxiv, 2020: p. 2020.12.03.408567.

60. Andersson, J.L., S. Skare, and J. Ashburner, How to correct susceptibility distortions in spin-echo echo-planar images: application to diffusion tensor imaging. Neuroimage, 2003. 20(2): p. 870–88.

61. Glasser, M.F., et al., The minimal preprocessing pipelines for the Human Connectome Project. Neuroimage, 2013. 80: p. 105–24.

62. Cai, L.Y., et al., PreQual: An automated pipeline for integrated preprocessing and quality assurance of diffusion weighted MRI images. Magnetic Resonance in Medicine, 2021. n/a(n/a).

63. Jenkinson, M., et al., Fsl. Neuroimage, 2012. 62(2): p. 782–90.

64. Wasserthal, J., et al., Combined tract segmentation and orientation mapping for bundle-specific tractography. Med Image Anal, 2019. 58: p. 101559.

65. Wasserthal, J., P. Neher, and K.H. Maier-Hein, TractSeg -Fast and accurate white matter tract segmentation. Neuroimage, 2018. 183: p. 239–253.

66. Wasserthal, J., P.F. Neher, and K.H. Maier-Hein. Tract Orientation Mapping for Bundle-Specific Tractography. in Medical Image Computing and Computer Assisted Intervention – MICCAI 2018. 2018. Cham: Springer International Publishing.

67. Tournier, J.D., et al., MRtrix3: A fast, flexible and open software framework for medical image processing and visualisation. Neuroimage, 2019. 202: p. 116137.

68. Yeh, F.-C., Shape analysis of the human association pathways. NeuroImage, 2020. 223: p. 117329.

69. Yeh, F.C., V.J. Wedeen, and W.Y. Tseng, Generalized q-sampling imaging. IEEE Trans Med Imaging, 2010. 29(9): p. 1626–35.

70. Yeh, F.C., et al., Population-averaged atlas of the macroscale human structural connectome and its network topology. Neuroimage, 2018. 178: p. 57–68.

71. Garyfallidis, E., et al., Recognition of white matter bundles using local and global streamline-based registration and clustering. Neuroimage, 2018. 170: p. 283–295.

72. Garyfallidis, E., et al., Dipy, a library for the analysis of diffusion MRI data. Front Neuroinform, 2014. 8: p. 8.

73. Warrington, S., et al., XTRACT - Standardised protocols for automated tractography in the human and macaque brain. Neuroimage, 2020. 217: p. 116923.

74. Tax, C.M.W., et al., The dot-compartment revealed? Diffusion MRI with ultra-strong gradients and spherical tensor encoding in the living human brain. Neuroimage, 2020. 210: p. 116534.

75. De Luca, A., et al., On the generalizability of diffusion MRI signal representations across acquisition parameters, sequences and tissue types: chronicles of the MEMENTO challenge. bioRxiv, 2021: p. 2021.03.02.433228.

76. Jones, D.K., M.A. Horsfield, and A. Simmons, Optimal strategies for measuring diffusion in anisotropic systems by magnetic resonance imaging. Magn Reson Med, 1999. 42(3): p. 515–25.

77. Chen, G., et al., Improving Estimation of Fiber Orientations in Diffusion MRI Using Inter-Subject Information Sharing. Scientific Reports, 2016. 6(1): p. 37847.

78. Moyer, D., et al., Scanner invariant representations for diffusion MRI harmonization. Magnetic Resonance in Medicine, 2020. 84(4): p. 2174–2189.

79. Huynh, K.M., et al. Longitudinal Harmonization for Improving Tractography in Baby Diffusion MRI. in Computational Diffusion MRI. 2019. Cham: Springer International Publishing.

80. Rheault, F., et al., Common misconceptions, hidden biases and modern challenges of dMRI tractography. J Neural Eng, 2020. 17(1): p. 011001.

81. Schilling, K., et al., Can increased spatial resolution solve the crossing fiber problem for diffusion MRI? NMR Biomed, 2017. 30(12).

82. Griswold, M.A., et al., Generalized autocalibrating partially parallel acquisitions (GRAPPA). Magn Reson Med, 2002. 47(6): p. 1202–10.

83. Pruessmann, K.P., et al., SENSE: sensitivity encoding for fast MRI. Magn Reson Med, 1999. 42(5): p. 952–62.

84. Prohl, A.K., et al., Reproducibility of Structural and Diffusion Tensor Imaging in the TACERN Multi-Center Study. Front Integr Neurosci, 2019. 13: p. 24.

85. Magnotta, V.A., et al., Multicenter reliability of diffusion tensor imaging. Brain Connect, 2012. 2(6): p. 345–55.

86. Min, J., et al., Inter-Vendor and Inter-Session Reliability of Diffusion Tensor Imaging: Implications for Multicenter Clinical Imaging Studies. Korean J Radiol, 2018. 19(4): p. 777–782.

87. Schilling, K.G., et al., Histological validation of diffusion MRI fiber orientation distributions and dispersion. Neuroimage, 2018. 165: p. 200–221.

88. Canales-Rodriguez, E.J., et al., Sparse wars: A survey and comparative study of spherical deconvolution algorithms for diffusion MRI. Neuroimage, 2018. 184: p. 140–160.

89. Tournier, J.D., F. Calamante, and A. Connelly, Determination of the appropriate b value and number of gradient directions for high-angular-resolution diffusion-weighted imaging. NMR Biomed, 2013. 26(12): p. 1775–86.

90. Tournier, J.D., et al., Resolving crossing fibres using constrained spherical deconvolution: validation using diffusion-weighted imaging phantom data. Neuroimage, 2008. 42(2): p. 617–25.

91. Prckovska, V., et al., Optimal acquisition schemes in high angular resolution diffusion weighted imaging. Med Image Comput Comput Assist Interv, 2008. 11(Pt 2): p. 9–17.

92. Daducci, A., et al., Quantitative comparison of reconstruction methods for intra-voxel fiber recovery from diffusion MRI. IEEE Trans Med Imaging, 2014. 33(2): p. 384–99.

93. Yeh, F.C., et al., Deterministic diffusion fiber tracking improved by quantitative anisotropy. PLoS One, 2013. 8(11): p. e80713.

94. Sotiropoulos, S.N., T.E. Behrens, and S. Jbabdi, Ball and rackets: Inferring fiber fanning from diffusion-weighted MRI. Neuroimage, 2012. 60(2): p. 1412–25.

95. Mandonnet, E., S. Sarubbo, and L. Petit, The Nomenclature of Human White Matter Association Pathways: Proposal for a Systematic Taxonomic Anatomical Classification. Front Neuroanat, 2018. 12: p. 94.

96. Smith, S.M., et al., Tract-based spatial statistics: Voxelwise analysis of multi-subject diffusion data. NeuroImage, 2006. 31(4): p. 1487–1505.

97. Vanderweyen, D.C., et al., The role of diffusion tractography in refining glial tumor resection. Brain Structure and Function, 2020. 225(4): p. 1413–1436.

98. Mancini, M., et al., Automated fiber tract reconstruction for surgery planning: Extensive validation in language-related white matter tracts. Neuroimage Clin, 2019. 23: p. 101883.

99. Fekonja, L., et al., Manual for clinical language tractography. Acta Neurochirurgica, 2019. 161(6): p. 1125–1137.

100. Essayed, W.I., et al., White matter tractography for neurosurgical planning: A topography-based review of the current state of the art. Neuroimage Clin, 2017. 15: p. 659–672.

101. Koller, K., et al., MICRA: Microstructural image compilation with repeated acquisitions. NeuroImage, 2021. 225: p. 117406.

102. Avesani, P., et al., The open diffusion data derivatives, brain data upcycling via integrated publishing of derivatives and reproducible open cloud services. Scientific Data, 2019. 6(1): p. 69.

103. Jack, C.R., Jr., et al., The Alzheimer’s Disease Neuroimaging Initiative (ADNI): MRI methods. J Magn Reson Imaging, 2008. 27(4): p. 685–91.

104. Chandio, B.Q., et al., Bundle analytics, a computational framework for investigating the shapes and profiles of brain pathways across populations. Sci Rep, 2020. 10(1): p. 17149.

105. Tong, Q., et al., Multicenter dataset of multi-shell diffusion MRI in healthy traveling adults with identical settings. Sci Data, 2020. 7(1): p. 157.

106. Galluzzi, S., et al., Clinical and biomarker profiling of prodromal Alzheimer’s disease in workpackage 5 of the Innovative Medicines Initiative PharmaCog project: a ‘European ADNI study’. J Intern Med, 2016. 279(6): p. 576–91.

107. Marcus, D.S., et al., Open Access Series of Imaging Studies (OASIS): cross-sectional MRI data in young, middle aged, nondemented, and demented older adults. J Cogn Neurosci, 2007. 19(9): p. 1498–507.

108. Zhang, F., et al., Deep white matter analysis (DeepWMA): Fast and consistent tractography segmentation. Med Image Anal, 2020. 65: p. 101761.

109. Zhang, F., et al., SlicerDMRI: Diffusion MRI and Tractography Research Software for Brain Cancer Surgery Planning and Visualization. JCO Clin Cancer Inform, 2020. 4: p. 299–309.

110. Zhang, F., et al., Deep white matter analysis: fast, consistent tractography segmentation across populations and dMRI acquisitions. Med Image Comput Comput Assist Interv, 2019. 11766: p. 599–608.

111. O’Donnell, L.J. and C.F. Westin, Automatic tractography segmentation using a high-dimensional white matter atlas. IEEE Trans Med Imaging, 2007. 26(11): p. 1562–75.

112. Wassermann, D., et al., The white matter query language: a novel approach for describing human white matter anatomy. Brain Struct Funct, 2016. 221(9): p. 4705–4721.

113. Ros, C., et al., Atlas-guided cluster analysis of large tractography datasets. PLoS One, 2013. 8(12): p. e83847.

114. Zöllei, L., et al., TRActs constrained by UnderLying INfant anatomy (TRACULInA): An automated probabilistic tractography tool with anatomical priors for use in the newborn brain. Neuroimage, 2019. 199: p. 1–17.

115. Schilling, K.G., et al., Tractography dissection variability: what happens when 42 groups dissect 14 white matter bundles on the same dataset? bioRxiv, 2021: p. 2020.10.07.321083.

116. Ning, L., et al., Cross-scanner and cross-protocol multi-shell diffusion MRI data harmonization: Algorithms and results. Neuroimage, 2020. 221: p. 117128.

117. Tax, C.M., et al., Cross-scanner and cross-protocol diffusion MRI data harmonisation: A benchmark database and evaluation of algorithms. Neuroimage, 2019.

